# Fat-rich diet reprograms intrapulmonary neutrophils to boost tissue-specific antitumor immunity

**DOI:** 10.1101/2025.05.20.655047

**Authors:** Yanling Wang, Jinjing Zhang, Ying Li, Tao Wang, Lu Wang, Yuxuan Miao, Chia-Wei Chang, Yuanyuan Liu, Ziyang Huang, Zhiqiu Yao, He Xu, Quan D. Zhou, Lei Wang, Yushi Yao

## Abstract

Neutrophils adapt to tissue-specific signals and exert innate defense functions against infections and malignancies. Fat-rich diet (FRD), such as high-fat diet (HFD) and ketogenic diet (KD), has complex impacts on immunity. However, whether and how FRD shapes tissue-specific functions of neutrophils remain unclear. Here we show that both isocaloric HFD- and KD-fed mice demonstrate enhanced neutrophil-mediated pulmonary tumor resistance than chow diet-fed mice. Intrapulmonary but not systemic neutrophils in FRD-fed mice bear enhanced potential of reactive oxygen species (ROS) production and ROS-dependent tumor cytotoxicity. Mechanistically, FRD-induced increased serum saturated fatty acids and cholesterol stimulate lung vascular endothelial cells (LVECs), which reprogram intrapulmonary neutrophils via contact- and intercellular adhesion molecule-1 (ICAM-1)-dependent mechanisms. Analysis on human lung single-cell RNA sequencing data showed that intensified cell adhesion and priming signals from human LVECs are associated with enhanced antitumor functions in intrapulmonary neutrophils. Our findings highlight the roles of dietary fats in shaping neutrophil functions in a tissue-specific manner. Dietary intervention targeting tissue-specific reprogramming of neutrophils therefore represents a potential strategy against malignancies in the lungs.

## Introduction

As one of the most abundant circulating leukocytes, neutrophils are quick-responding innate myeloid cells which play critical roles in anti-infectious, antitumor, and autoimmune responses ^1^. Functional modulation of neutrophils in both the bone marrow and peripheral tissues is therefore highly relevant to host susceptibility to various diseases ^1–13^. In the bone marrow, reprogrammed granulopoiesis of neutrophils was shown to induce systemically altered neutrophil functions ^14, 15^. In peripheral tissues, despite the short half-life of neutrophils in the circulation of typically less than one day, emerging evidence has just started to uncover their previously unexpected degree of functional diversification induced by tissue-specific factors such as those in the lung, intestine, and skin ^9, 16^.

Recent studies highlight the lung vasculature as a key place where tissue-specific functional reprogramming of neutrophils occurs ^9, 16, 17^. Specifically, lung vascular endothelial cells (LVECs) guide steady-state intrapulmonary neutrophils to vascular niches where neutrophils are reprogrammed in a tissue-specific manner to support vascular growth ^16^. Despite these intriguing findings, the functional significance and immunological mechanisms of tissue-specific neutrophil reprogramming in pathological scenarios remain not well understood ^5, 7^. With the lung a preferential target of both primary and metastatic malignancies ^18, 19^, it remains unclear whether and how pulmonary tissue-specific signals shape antitumor functions in neutrophils.

Dietary fat contents have profound and complex influences on clinical outcomes of various diseases, and dietary intervention by changing fat contents has long been considered for potential clinical application in patients with various diseases including cancers ^20–26^. Although over intake of dietary fats for extended time typically induces gain of fat tissue mass and obesity in both human and experimental animals, accumulating evidence has suggested that the impact of dietary fat contents on the immune system can well be independent on and drastically different from that of obesity ^3, 8, 20, 22, 27–29^. In this regard, little is known on whether and how dietary fat contents influence antitumor functions of neutrophils independently of excessive calorie intake and obesity.

To determine the impact of dietary fat contents on tissue-specific reprograming of neutrophils in the lung, we inoculated either metastatic or primary lung tumor cells into the lungs of mice fed with either a non-obese isocaloric HFD strategy or KD. We show here that FRD (i.e., isocaloric HFD and KD) induces tissue-specific antitumor responses in the lung, which is mediated by intrapulmonary neutrophils reprogrammed by FRD-exposed LVECs. Our study therefore reveals a previously unappreciated role of dietary fat contents in shaping tissue-specific antitumor functions of neutrophils in the lung.

## Results

### FRD induces antitumor response in the lung

To determine the impact of FRD on pulmonary antitumor responses, wild-type C57BL/6 mice were fed with either chow diet (CD) or HFD (*ad libitum* through experimental endpoint) for 4 weeks, followed by injection of luciferase-expressing B16F10 melanoma cells (B16-luc) intravenously (i.v.) to inoculate tumor cells in the lungs (Fig. 1a) ^30^. Such a short-term HFD-feeding strategy is isocaloric, as demonstrated by comparable average daily calorie intake in HFD- and CD-fed mice (Extended Data Fig. 1a-e), as well as comparable body weight in both groups of mice through the endpoint at 6 weeks after the initiation of HFD (Extended Data Fig. 1f). Compared to CD-fed mice, reduced luciferin tumor signals in the lung area were observed in HFD-fed mice at various time points after tumor inoculation (Fig. 1b,c). Consistent with *in vivo* tumor imaging observations, macroscopic evaluation of the lungs at the experimental endpoint showed reduced visible B16 tumor nodules in lung lobes of HFD than CD mice (Fig. 1d,e). Lung histopathological analysis also showed lower percentages of lung area occupied by B16 tumor lesions in HFD than in CD mice (Fig. 1f,g). To determine whether such an HFD-induced antitumor response is a tissue-specific or systemic phenotype, we alternatively injected B16 melanoma cells subcutaneously (s.c.) (Extended Data Fig. 2a). Analysis on tumor volume at the injection site at various time points and tumor weight at endpoint showed that s.c. B16 tumor growth was comparable in CD and HFD mice (Extended Data Fig. 2b-d), indicating that HFD does not enhance systemic antitumor response.

**Figure 1.**
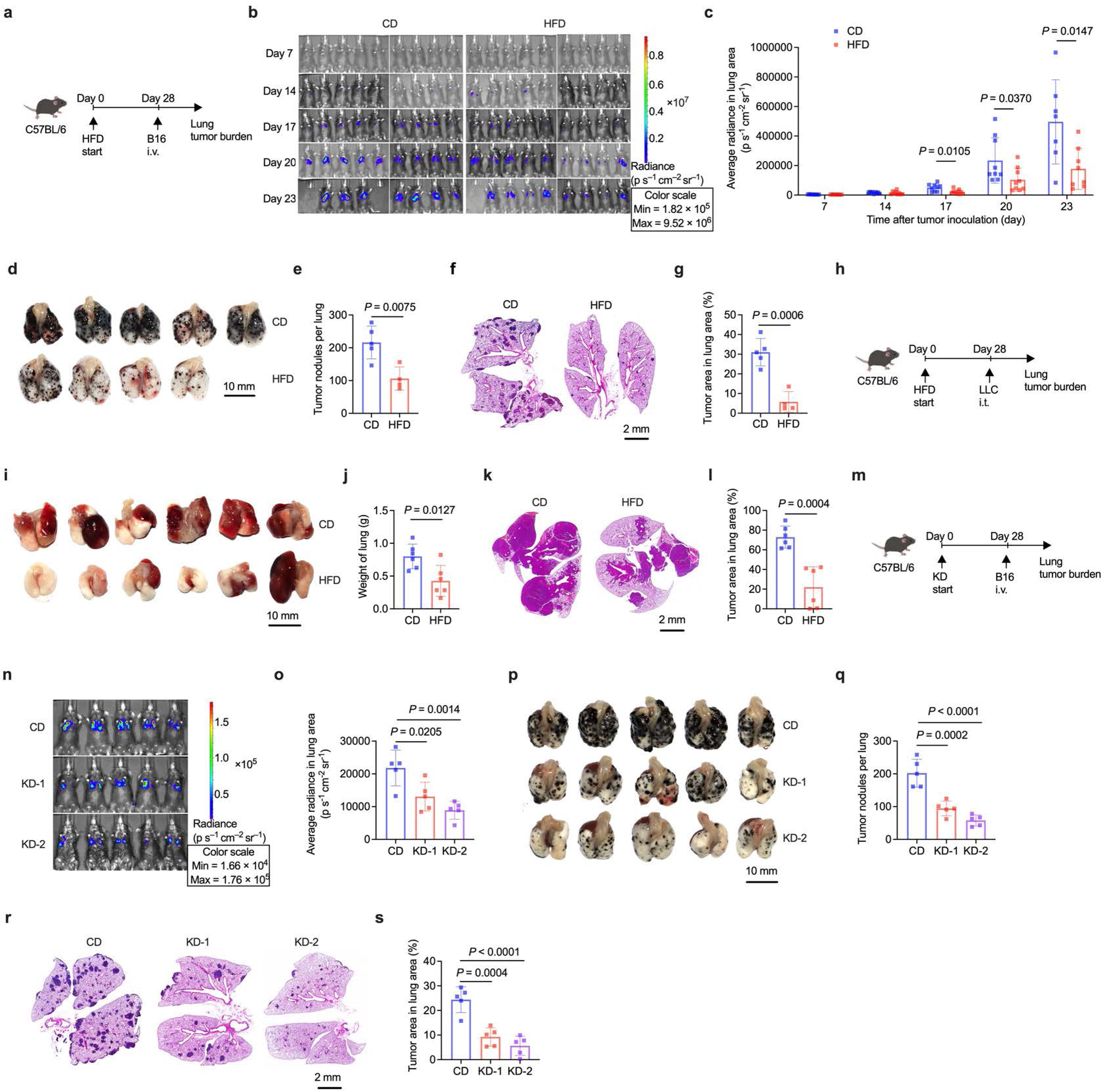
FRD induces antitumor response in the lung. (**a**) Schema of experimental design showing high-fat diet (HFD) or chow diet (CD) feeding schedule in C57BL/6 mice followed by intravenous (i.v.) injection of luciferase-expressing B16 melanoma cells. (**b**,**c**) Representative bioluminescence imaging (**b**) and quantification of average radiance of luciferin signals in the lung region (**c**) on days 7 to 23 after tumor inoculation in CD versus HFD mice. n = 9 mice per group. Deceased animals are indicated by vacancy in (**b**). (**d**,**e**) Representative macroscopic lung images (**d**) and quantification of visible B16 tumor nodules (**e**) on the surface of lung lobes at day 16 after tumor inoculation in CD versus HFD mice. n = 5 mice in CD group and n = 4 in HFD group. (**f**,**g**) Representative lung histopathology images (**f**) and percentage of lung area occupied by tumor lesions (**g**) based on lung histopathological analysis in mice shown in (**d**). (**h**) Schema of intratracheal (i.t.) Lewis lung carcinoma (LLC) cell inoculation in HFD or CD mice. (**i**,**j**) Representative macroscopic lung images (**i**) and weight of lung (**j**) at experimental endpoint. n = 6 mice per group. (**k**,**l**) Representative lung histopathology images (**k**) and percentage of lung area occupied by tumor lesions (**l**) based on lung histopathological analysis in mice shown in (**i**). (**m**) Schema of experimental design showing ketogenic diets (KD-1: 90% lard-derived fat; KD-2: 90% plant oil-derived fat) feeding schedule in C57BL/6 mice followed by intravenous (i.v.) injection of luciferase-expressing B16 melanoma cells. (**n**,**o**) Representative bioluminescence imaging (**n**) and quantification of average radiance of luciferin signals in the lung region (**o**) at day 16 after tumor inoculation in CD versus KD mice. n = 5 mice per group. (**p**,**q**) Representative macroscopic lung images (**p**) and quantification of visible tumor nodules (**q**) on the surface of lung lobes at day 16 after tumor inoculation in mice shown in n. (**r**,**s**) Representative lung histopathology images (**r**) and percentage of lung area occupied by tumor lesions (**s**) based on lung histopathological analysis in mice shown in (**n**). Data are representative of two (**b**,**c**,**i**-**s**) or three (**d**-**g**) independent experiments. Bar graphs are presented as mean ± s.d. Two-tailed Student’s t-test was used for comparison between two-groups. One-way ANOVA followed by a Tukey test was performed to compare more than two groups.

To assess whether reduced lung B16 tumor burden in HFD mice is due to reduced initial tumor extravasation and inoculation in the lung, we quantified tumor cells in lung tissues at 24 h after i.v. injection of B16-luc cells expressing green fluorescent protein (GFP). By quantifying CD45^−^GFP^+^ B16 tumor cells and B16 cell-specific gene transcripts of dopachrome tautomerase (*Dct*), premelanosome (*Pmel*), and luciferase ^30^, we observed comparable early tumor burdens in lung tissues between CD and HFD mice (Extended Data Fig. 3a,b). These data suggest that HFD-induced pulmonary antitumor response is not due to reduced initial inoculation of B16 tumor cells to the lung tissues. We also assessed the impact of HFD on pulmonary antitumor response in a mouse model of primary lung cancer where LLC lung cancer cells were inoculated via intratracheal (i.t.) distillation (Fig. 1h). Consistent with our findings in B16 i.v. lung tumor model, we observed reduced weight of LLC tumor-bearing lungs (Fig. 1i,j) and lower percentages of lung area occupied by LLC tumor lesions in HFD than CD mice (Fig. 1k,l). These data thus suggest that HFD induces antitumor response in the lung.

We next sought to determine whether HFD-induced pulmonary antitumor phenotype requires such a strict four-week HFD feeding prior to tumor inoculation. We therefore fed mice with HFD for 8 weeks before inoculation of B16 melanoma cells i.v. (Extended Data Fig. 4a). Pulmonary antitumor response was consistently observed in mice fed in such a prolonged HFD formula, as evidenced by reduced lung tumor burdens both macroscopically and microscopically at endpoint (Extended Data Fig. 4b-e). We could also observe pulmonary antitumor phenotype in mice fed with HFD for 4 weeks prior to B16 i.v., while switching to CD immediately after tumor inoculation (Extended Data Fig. 4f-j), as well as in mice with HFD feeding only immediately after B16 i.v. (Extended Data Fig. 4k-o). When we inoculated B16 melanoma cells i.v. in mice previously fed with HFD for 4 weeks and switched to CD for another 4 weeks before tumor inoculation, we observed moderate antitumor response in the lung, as evidenced by moderately reduced B16 tumor nodules in lung lobes and significantly reduced percentage of lung area occupied by tumor lesions (Extended Data Fig. 4p-t). These data collectively suggest that HFD induces pulmonary antitumor response regardless of whether HFD is introduced before or after tumor inoculation, and HFD-induced antitumor response is not long-lasting after discontinuation of HFD.

KD represents a FRD characterized by extremely low carbohydrates ^21, 25^. We therefore tested whether KD could also induce antitumor response in the lung. To do this, mice were fed with CD or KD (i.e., either a KD-1 formula with 90% lard-derived fat, or a KD-2 with 90% plant oil) for 4 weeks, followed by inoculation of B16-luc melanoma cells i.v. (Fig. 1m). Similar to HFD mice, we observed reduced lung tumor burdens in mice fed with either KD formula, as evidenced by reduced luciferin tumor signals in the lung area (Fig. 1n,o), reduced visible B16 tumor nodules in lung lobes (Fig. 1p,q), and lower percentages of lung area occupied by B16 tumor lesions in lung histopathological analysis (Fig. 1r,s). These data collectively suggest that FRD, including isocaloric HFD and KD, induces antitumor response in the lung in a tissue-specific manner.

### FRD-induced pulmonary antitumor response is dependent on neutrophils

We next sought to determine whether FRD-induced pulmonary antitumor phenotype is dependent on boosted antitumor immunity and, if so, which leukocyte subset is primarily involved. Alveolar macrophages (AMs) play critical roles in innate antitumor immunity in the lung ^30^. We therefore depleted AMs immediately before and after B16 melanoma cell inoculation (i.v.) in CD or HFD mice (Extended Data Fig. 5a). Although depletion of AMs resulted in increased lung tumor burdens in both CD and HFD mice, AM-depleted HFD mice still had reduced lung tumor burdens as compared to their CD counterparts (Extended Data Fig. 5b-e), indicating that HFD-induced antitumor response is independent of AMs. To directly compare the antitumor potential of AMs, we purified AMs from CD and HFD mice and coculture them respectively with B16-luc cells to determine tumor cytotoxic functions of AMs ^30^. After 48 hours of coculture, we observed comparable tumor cell survival in B16 tumor cells cocultured with CD AMs and those cocultured with HFD AMs (Extended Data Fig. 5f). These data suggest that HFD does not alter innate antitumor functions of AMs. To determine whether natural killer (NK) cells and T cells might contribute to HFD-induced antitumor response in the lung, we depleted either NK cells or CD4^+^ and CD8^+^ T cells *in vivo* before and after B16 melanoma cell inoculation (i.v.) in CD or HFD mice (Extended Data Fig. 5g). Although depletion of NKs resulted in evidently increased lung tumor burdens in both CD and HFD mice ^31^, NK cell-depleted HFD mice showed reduced lung tumor burdens as compared to their CD counterparts (Extended Data Fig. 5h-k). Depletion of both CD4^+^ and CD8^+^ T cells did not alter the reduced lung tumor burdens in HFD mice (Extended Data Fig. 5h-k). These data suggest that neither NK cells nor T cells are required for HFD-induced antitumor response in the lung.

Neutrophils play critical roles in anti-tumor immunity ^6, 32^. Indeed, by using an intravascular staining strategy ^33^, we found that neutrophils resided exclusively in lung vasculature before tumor inoculation in both CD and HFD mice, and they infiltrated extravascular lung tissues after tumor inoculation and were found in lung tumor lesions (Fig. 2a,b). To determine whether neutrophils are required for HFD-induced antitumor response, we depleted neutrophils *in vivo* in wild-type mice by repeated administration of anti-Ly6G depleting antibodies immediately before and after B16 melanoma cell inoculation (i.v.) in CD or HFD mice (Fig. 2c and Extended Data Fig. 6a-c). Neutrophil depletion did not alter lung tumor burdens in CD mice (Fig. 2d-i). In contrast, neutrophil depletion resulted in increased lung tumor burdens in HFD mice to levels comparable to CD mice (Fig. 2d-i), suggesting a critical role of neutrophils in HFD-induced pulmonary antitumor immunity. In a similar experimental setting, we administered repeated doses of diphtheria toxin (DT) to deplete neutrophils in CD- or HFD-fed Ly6G-DTR mice, followed by i.v. inoculation of B16 melanoma cells (Fig. 2j and Extended Data Fig. 6d). We consistently observed that DT-mediated neutrophil depletion in Ly6G-DTR mice abrogated HFD-induced pulmonary antitumor immunity (Fig. 2k-n). To test if neutrophils are also required for KD-induced pulmonary antitumor immunity, we depleted neutrophils in CD or KD Ly6G-DTR mice by repeated doses of DT, followed by i.v. inoculation of B16 melanoma cells (Fig. 2o). We observed abrogated anti-tumor response in neutrophil-depleted KD mice (Fig. 2p-s). These data suggest that FRD-induced pulmonary antitumor immunity is dependent on neutrophils.

**Figure 2.**
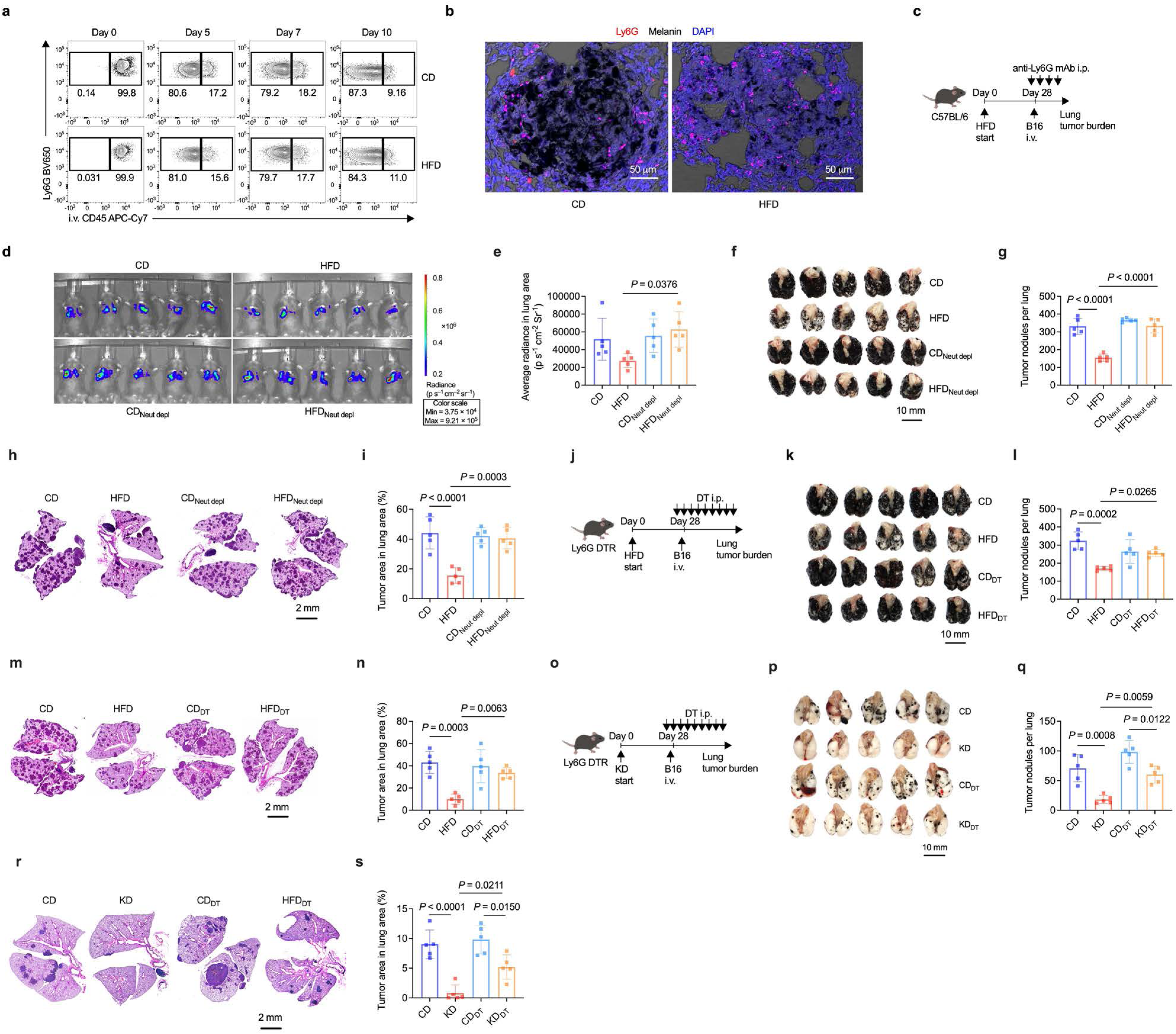
FRD-induced pulmonary antitumor response is dependent on neutrophils. (a) Representative flow cytometry contour plots of lung neutrophil populations using intravascular (i.v.) staining with anti-CD45 antibody at before (day 0) and 5, 7, and 10 days after B16-luc melanoma cell inoculation in HFD and CD mice. Numbers beside gates represent percentages in parental gates. (b) Representative confocal microscopic images on Ly6G^+^ neutrophils in lung B16 tumor lesions in CD and HFD mice. (c) Schema of experimental design showing continuous neutrophil depletion in *vivo* by repeated doses of anti-Ly6G antibodies in HFD and CD mice immediately before and after i.v. inoculation of B16-luc melanoma cells. (**d**,**e**) Representative bioluminescence imaging (**d**) and quantification of average radiance of luciferin signals in the lung region (**e**) at day 16 after tumor inoculation. n = 5 mice per group. (**f**,**g**) Representative macroscopic lung images (**f**) and quantification of visible tumor nodules (**g**) on the surface of lung lobes in mice shown in (**d**). (**h**,**i**) Representative lung histopathology images (**h**) and percentage of lung area occupied by tumor lesions (**i**) based on lung histopathological analysis in mice shown in (**d**). (**j**) Schema of experimental design of diphtheria toxin (DT)-mediated neutrophil depletion in HFD- or CD-fed Ly6G-DTR mice immediately before and after i.v. inoculation of B16 melanoma cells. (**k**,**l**) Representative macroscopic lung images (**k**) and quantification of visible tumor nodules (**l**) on the surface of lung lobes. n = 5 mice per group. (**m**,**n**) Representative lung histopathology images (**m**) and percentage of lung area occupied by tumor lesions (**n**) based on lung histopathological analysis in mice shown in (**k**). (**o**) Schema of experimental design of DT-mediated neutrophil depletion in KD (90% lard-derived fat)- or CD-fed Ly6G-DTR mice immediately before and after i.v. inoculation of B16 melanoma cells. (**p**,**q**) Representative macroscopic lung images (**p**) and quantification of visible tumor nodules (**q**) on the surface of lung lobes. n = 5 mice per group. (**r**,**s**) Representative lung histopathology images (**r**) and percentage of lung area occupied by tumor lesions (**s**) based on lung histopathological analysis in mice shown in (**p**). Data are representative of two (**a**,**b**,**d**-**i**) or three (**k**-**s**) independent experiments. Bar graphs are presented as mean ± s.d. One-way ANOVA followed by a Tukey test was performed to compare multiple groups.

### FRD boosts tumor-cytotoxic functions in lung neutrophils

Our data thus far suggest that FRD induces lung tissue-specific antitumor immunity in a neutrophil-dependent manner. We therefore speculate that neutrophils in the lung tissues of FRD-fed mice have enhanced antitumor functions. To characterize FRD-exposed lung neutrophils, we performed transcriptional, phenotypic, and functional analysis in neutrophils isolated from lung tissues of CD and HFD mice (Fig. 3a). RNA sequencing (RNA-seq) in lung neutrophils showed that the transcription of 528 genes upregulated and 466 downregulated in HFD versus CD lung neutrophils (Fig. 3b). Moreover, gene ontology (GO) enrichment of cluster-specific marker genes revealed upregulation of gene transcripts associated with immune activation and effector functions in HFD versus CD lung neutrophils (Fig. 3c). Notably, HFD lung neutrophils had enhanced transcription of gene clusters related to cell killing and ROS biosynthetic process that are closely related to their innate antitumor functions (Fig. 3d). In addition to transcriptional changes, flow cytometry analysis showed that HFD lung neutrophils expressed higher levels of CD177, CD11b, Ly6G, and had higher autofluorescence than their CD counterparts (Fig. 3e) ^16, 32, 34^. These data suggest that HFD induces transcriptional and phenotypic changes in lung neutrophils.

**Figure 3.**
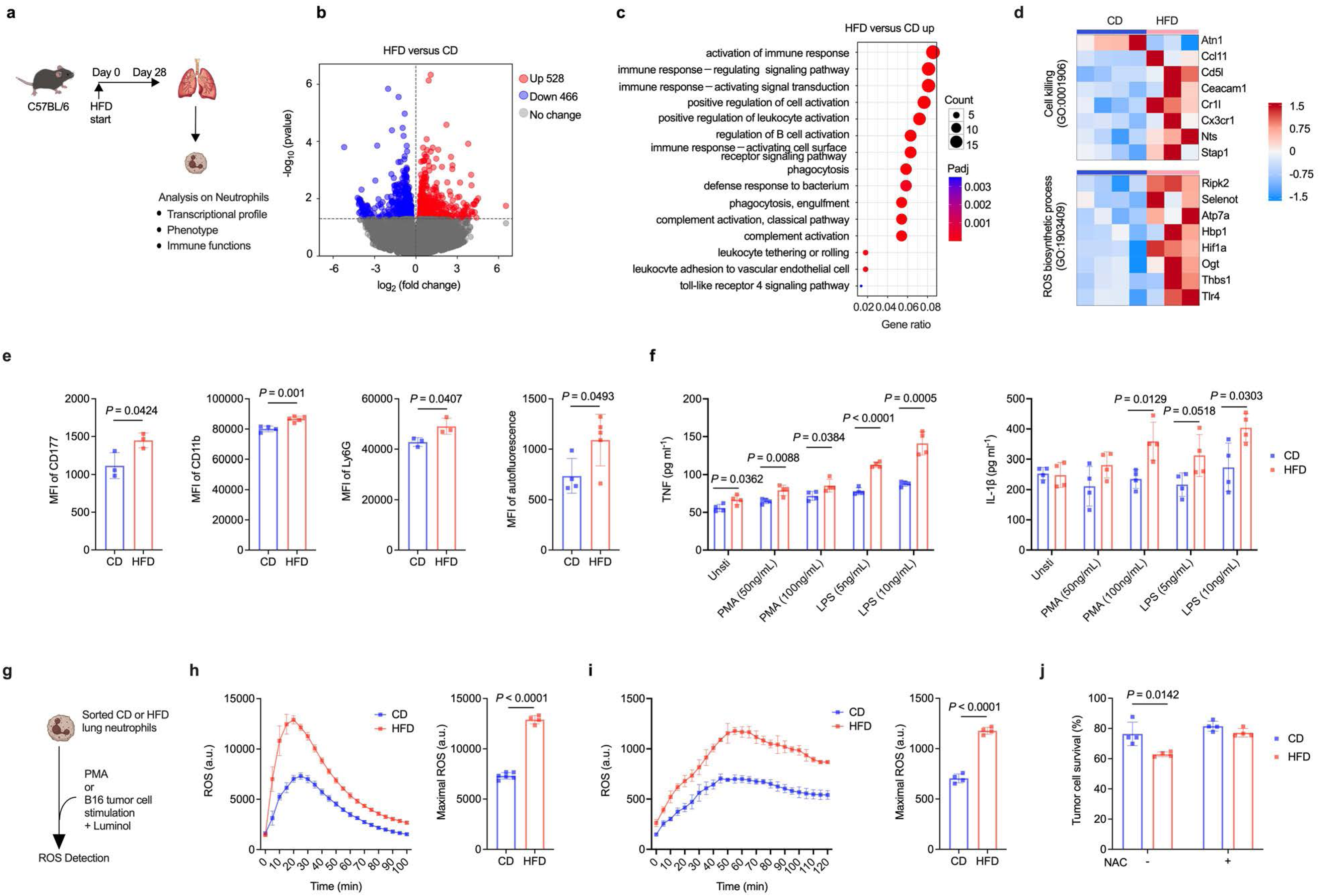
FRD boosts tumor-cytotoxic functions in lung neutrophils. (a) Schema of experimental design of transcriptional, phenotypic, and functional analysis of lung neutrophils at 28 days after initiation of HFD or CD feeding. (b) Volcano plot comparing transcriptomes of lung neutrophils from HFD versus CD mice (DESeq2, P < 0.05, |log2(fold change) | > 0). n = 4 in CD group and n = 3 in HFD group. (c) Gene Ontology (GO) enrichment analysis of significantly upregulated gene transcripts in lung neutrophils from HFD versus CD mice (P < 0.05). n = 4 in CD group and n = 3 in HFD group. (d) Heatmaps of DEGs (z-score normalized, P < 0.05) related to cell killing and ROS biosynthetic processes in lung neutrophils from HFD and CD mice. n = 4 in CD group and n = 3 in HFD group. (e) Median fluorescence intensity (MFI) of selected surface markers differentially expressed on lung neutrophils in HFD versus CD mice. n = 3 or 4 mice per group as indicated. (f) Contents of TNF and IL-1β in culture supernatants of *ex vivo* cultured lung neutrophils stimulated with PMA or LPS. n = 4 culture wells per group. (g) Schema of quantification of ROS in lung neutrophils stimulated with PMA or B16 melanoma cells. (h) ROS contents in culture supernatants of PMA-stimulated lung neutrophils. Left: kinetics of ROS contents in arbitrary units (a.u.). Right: peak ROS levels. n = 6 culture wells in CD group and n = 4 in HFD group. (i) ROS contents in culture supernatants of B16 cell-stimulated lung neutrophils. Left: kinetics of ROS contents in arbitrary units (a.u.). Right: peak ROS levels. n = 4 culture wells per group. (j) Percentage of B16 melanoma cell survival after coculture with CD or HFD lung neutrophils at an effector to target ratio (E:T) of 10:1, with or without supplementation of N-Acetyl-Cysteine (NAC; 5mM) to cell culture media. Data are representative of two (**f**,**i**,**j**) or three (**e**,**h**) independent experiments, or from one experiment (**b**-**d**). Graphs in (**e**,**f**,**h**-**j**) are presented as mean ± s.d. Two-tailed Student’s t-test was performed for comparison between two groups.

To assess functional differences between CD and HFD lung neutrophils, we purified neutrophils from the lungs of CD or HFD mice and stimulated them *ex vivo* with Phorbol-12-myristate-13-acetate (PMA) or lipopolysaccharide (LPS), followed by quantification of proinflammatory cytokines in the culture supernatants at 12 hours after stimulation. We detected higher amounts of tumor necrosis factor (TNF) and interleukin-1 beta (IL-1β) in the supernatants of HFD than CD neutrophils (Fig. 3f), suggesting enhanced immune-reactivity of HFD lung neutrophils. We also quantified ROS in the supernatant of neutrophils stimulated *ex vivo* with either PMA or B16 tumor cells. Notably, HFD lung neutrophils produced a higher amount of ROS in response to both PMA and B16 melanoma cells, as compared to CD neutrophils (Fig 3g-i), further suggesting their enhanced responsiveness related to antitumor functions. To directly assess the antitumor functions of CD versus HFD lung neutrophils, we cocultured B16-luc cells with freshly isolated CD or HFD lung neutrophils. After 48 hours of coculture, there were significantly reduced survival of B16 cells cocultured with HFD lung neutrophils compared to those cocultured with CD ones (Fig. 3j). Importantly, such an increased tumor cytotoxic effect by HFD lung neutrophils was abrogated when N-acetyl-cysteine (NAC), a scavenger of ROS ^15^, was supplemented to cell culture media (Fig. 3j). These data suggest that HFD lung neutrophils have enhanced tumor cytotoxic functions via increased production of ROS in response to tumor cells.

### Neutrophils acquire enhanced antitumor functions locally in FRD-exposed lung

Functional reprogramming of neutrophils may occur either systemically (e.g., in bone marrow) or locally in peripheral non-hematopoietic tissues such as the lung ^9^. We therefore asked whether HFD-induced pulmonary antitumor immunity is due to systemic or tissue-specific reprogramming of neutrophils. To test whether functional changes in lung neutrophils from HFD mice result from systemic neutrophil reprogramming, we compared transcriptional profiles in HFD and CD lung neutrophils with those from peripheral blood (PB) and the spleen (Fig. 4a). There were small number of differentially expressed gene transcripts between CD and HFD PB neutrophils, with the transcription of 35 genes upregulated and 66 downregulated in HFD versus CD PB neutrophils (Fig. 4b). Transcriptional differences were more evident in HFD versus CD splenic neutrophils (Fig. 4c), thought differentially-expressed genes (DEGs) in HFD versus CD splenic neutrophils were largely irrelevant to those in lung neutrophils, with only 53 overlapping DEGs among 1362 DEGs in HFD vs CD splenic neutrophils and 994 DEGs in HFD vs CD lung neutrophils (Fig. 3b and Fig. 4d). These data thus indicate that transcriptional differences in lung neutrophils of HFD versus CD mice are not due to systemic transcriptional reprogramming of neutrophils, and therefore are likely induced locally in the lung.

**Figure 4.**
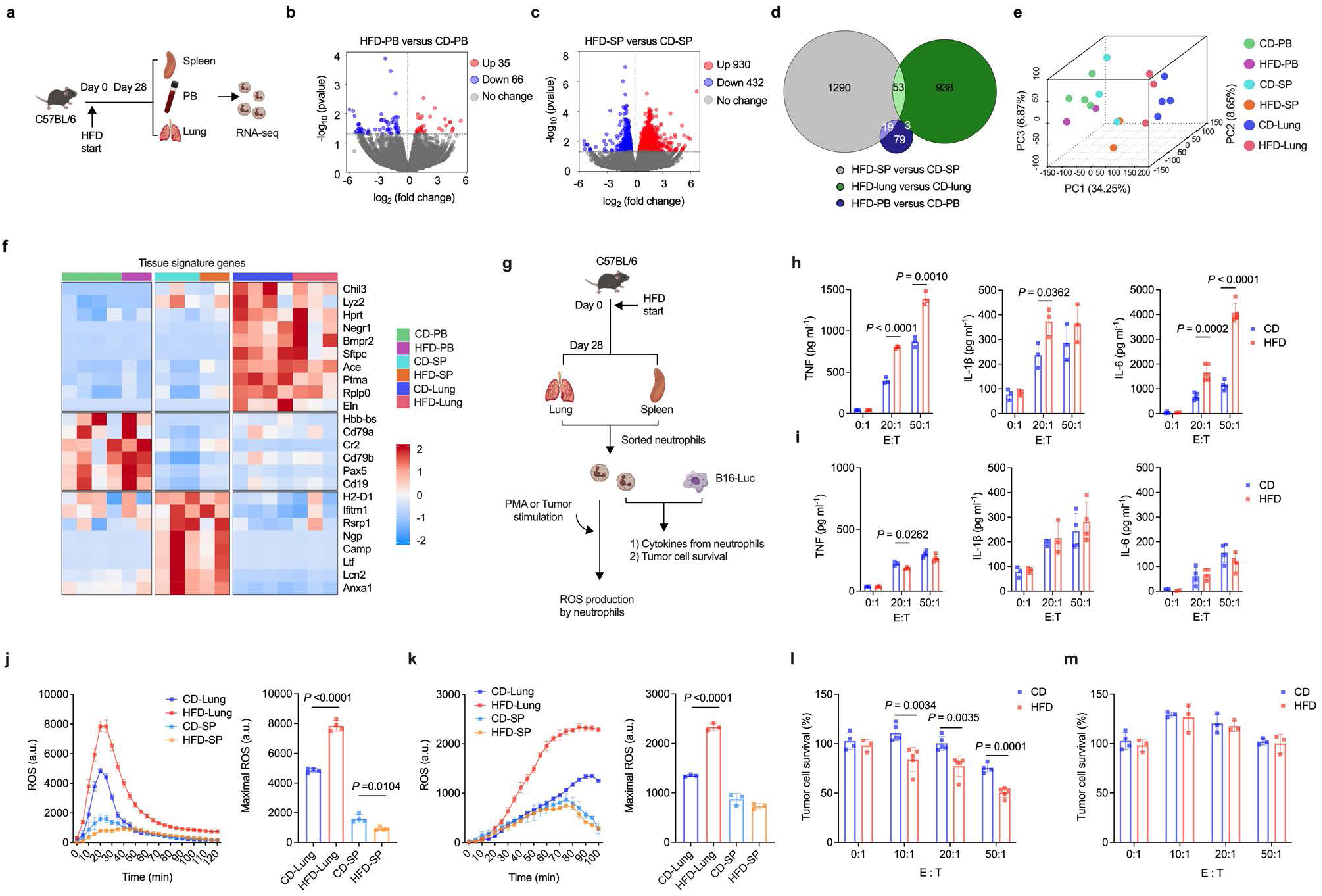
Neutrophils acquire enhanced antitumor functions locally in FRD-exposed lung. (a) Schema of transcriptional analysis in neutrophils from peripheral blood (PB), spleen (SP) and lung. (**b**,**c**) Volcano plot comparing transcriptomes of PB (**b**) and SP (**c**) neutrophils from HFD versus CD mice (DESeq2, P < 0.05, |log2(fold change)| > 0). (d) Venn diagram showing overlap of tissue-specific DEGs (P < 0.05) of HFD versus CD neutrophils. (e) 3D PCA of PB, SP, and lung neutrophils from CD or HFD mice. (f) Heatmaps showing DEGs of tissue-specific signature genes in PB, SP, and lung neutrophils (z-score normalized, P < 0.05). In (**b**-**f**), n = 4 in CD-PB and CD lung groups, n = 3 in CD-SP and HFD-lung groups, and n = 2 in HFD-PB and HFD-SP groups. (g) Schema of functional analysis in lung and SP neutrophils. (**h**,**i**) Contents of TNF, IL-1β and IL-6 in supernatants of lung (**h**) and spleen (**i**) neutrophils cocultured with B16 melanoma cells at various effector to target (E:T) ratios. n = 2, 3, 4 or 5 culture plates as indicated. (**j**,**k**) ROS contents in culture supernatants of PMA- (**j**) or B16 cell- (**k**) stimulated lung or SP neutrophils. Left: kinetics of ROS contents in arbitrary units (a.u.). Right: peak ROS levels. n = 3 or 4 culture wells per group as indicated. (**l**,**m**) Percentage of B16 melanoma cell survival after coculture with lung (**l**) or SP (**m**) neutrophils at various effector to target (E:T) ratios. n = 3, 4, or 5 culture wells per group as indicated. Data in (**h**-**m**) are representative of two independent experiments, and data in (**b**-**f**) are from one experiment. Graphs in (**h**-**m**) are presented as mean ± s.d. Two-tailed Student’s t-test was used for comparison between two-groups. One-way ANOVA followed by a Tukey test was performed to compare more than two groups.

Adaptation of neutrophils in the lung was recently shown to be a key factor in shaping tissue-specific neutrophil functions ^16^. Indeed, we observed a tissue-specific pattern of gene transcription in neutrophils from the spleen, PB, and lung, regardless of diet (Fig. 4e,f) ^16^. Specifically, there were drastic transcriptional changes in lung versus PB neutrophils (8874 DEGs in CD mice and 6340 in HFD mice, with 5877 DEGs overlapping) (Extended Data Fig. 7a-c), which were consistent with recent research findings showing tissue-specific transcriptional features of neutrophils in the lung ^16^. Transcriptional differences were also evident in lung versus splenic neutrophils in both CD and HFD mice (8710 DEGs in CD mice and 6110 in HFD mice, with 5472 DEGs overlapping) (Extended Data Fig. 7d-f). In contrast, transcriptional difference was less evident in splenic versus PB neutrophils (Extended Data Fig. 7g-i). In addition to transcriptional changes, we also observed differences in expression levels of cell surface molecules in lung versus splenic and PB neutrophils, in particular those associated with neutrophil functions such as CD11b, Ly6G, CD177, CD14, Siglec-F, CX3CR1, CD49D, and ICAM-1 (Extended Data Fig. 7j,k) ^16, 32, 34^. These data thus support an idea that enhanced lung neutrophil functions in HFD mice are acquired locally in lung tissues but not systemically.

To further show that HFD boosts immune functions in neutrophils specifically in the lung, we isolated lung or splenic neutrophils from CD or HFD mice, followed by stimulation of neutrophils *ex vivo* with B16 melanoma cells or PMA (Fig. 4g). After stimulation with B16 tumor cells, there were higher amount of proinflammatory cytokines, including TNF, IL-1β, and interleukin-6 (IL-6), in the supernatants of lung but not splenic neutrophils from HFD mice (Fig. 4h,i). Increased ROS contents were also observed in the culture supernatants of HFD lung but not splenic neutrophils stimulated with either PMA or tumor cells, as compared to their respective CD counterparts (Fig. 4j,k). More importantly, increased cytotoxic effect was observed in tumor cells cocultured with HFD than those cocultured with CD lung neutrophils (Fig. 4l). In contrast, splenic neutrophils from either CD or HFD mice did not show evident tumor cytotoxic functions (Fig. 4m). These data collectively suggest that neutrophils acquire enhanced tumor-cytotoxic functions locally in the lung of HFD mice.

### FRD-exposed LVECs boost neutrophil functions

Lung tissue-specific adaptation of neutrophils was shown to be guided by LVECs ^16^. Indeed, neutrophils reside exclusively in lung vasculature before tumor inoculation regardless of diet (Fig. 2a), we therefore speculate that LVECs play critical roles in FRD-induced innate antitumor immunity in lung neutrophils. We performed RNA-seq in lung neutrophils and observed increased gene transcripts related to cell-cell adhesion in neutrophils in the pre-tumoral lung tissues in HFD than CD mice (Fig. 5a), indicating intensified interactions between neutrophils and HFD-exposed LVECs. To characterize LVECs in HFD versus CD mice, we performed RNA-seq analysis in purified LVECs and found that the increased gene transcripts in HFD over CD LVECs are, although relatively small in number, mostly associated with immune functions including cell-cell adhesion and neutrophil chemotaxis (Fig. 5b-e). To further show the impact of HFD on LVECs, we performed intracellular flow cytometry staining on selected signal transduction molecules in LVECs, and observed increased contents of phosphorylated nuclear factor kappa-B (NF-κB) p-65 subunit (p-p65), phosphorylated mitogen-activated protein kinase (MAPK) p-38 subunit (p-p38), and phosphorylated extracellular regulated protein kinase (p-ERK) in LVECs from HFD than CD mice (Fig. 5f) ^35, 36^. These data suggest that HFD induces transcriptional changes and activation of intracellular signaling in LVECs.

**Figure 5.**
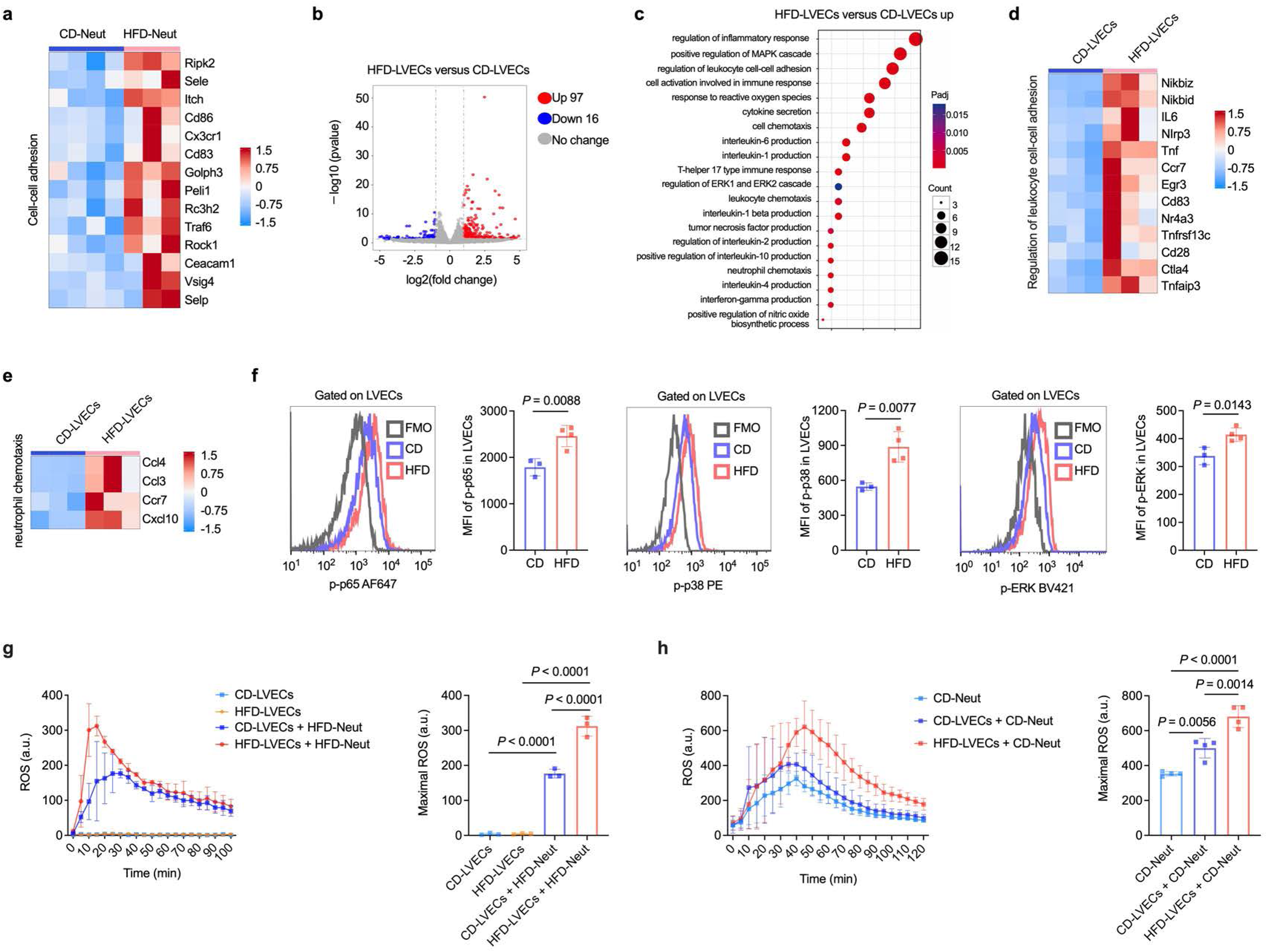
FRD-exposed LVECs boost neutrophil functions. (a) Heatmap showing DEGs related to cell-cell adhesion between HFD and CD lung neutrophils (z-score normalized, P < 0.05). n = 4 mice in CD group and n =3 mice in HFD group. (b) Volcano plot on transcriptional differences between lung vascular endothelial cells (LVECs) from HFD and CD mice (DESeq2, P < 0.05, |log2(fold change)| > 1). n = 3 mice per group in B-E. (c) GO enrichment analysis of significantly upregulated gene transcripts in LVECs from HFD versus CD mice (P < 0.05). (**d**,**e**) Heatmap of DEGs associated with regulation of leukocyte cell-cell adhesion (**d**) and neutrophil chemotaxis (**e**) in HFD versus CD LVECs (z-score normalized, P < 0.05). (f) Representative flow cytometry histograms and MFI of phosphorylated NF-κB p65 (p-p65), phosphorylated p38 MAPK (p-p38), and phosphorylated ERK (p-ERK) in HFD and CD LVECs. FMO: fluorescence minus one. n = 3 or 4 mice per group as indicated. (g) ROS contents in culture supernatants of PMA-stimulated HFD lung neutrophils cocultured overnight with CD or HFD LVECs prior to stimulation. Left: kinetics of ROS contents in arbitrary units (a.u.). Right: peak ROS levels. n = 3 culture wells per group. (h) ROS contents in culture supernatants of PMA-stimulated CD lung neutrophils cocultured overnight with CD or HFD LVECs prior to stimulation. Left: kinetics of ROS contents in arbitrary units (a.u.). Right: peak ROS levels. n = 4 culture wells per group. Data are representative of three (**f**-**g**) independent experiments, or from one experiment (**a**-**e**). Graphs with error bars are presented as mean ± s.d. Two-tailed Student’s t-test was used for comparison between two-groups. One-way ANOVA followed by a Tukey test was performed to compare more than two groups.

To directly address the impact of CD versus HFD LVECs on neutrophils, we isolated LVECs from CD or HFD mice and coculture them with isolated HFD lung neutrophils. After overnight coculture, neutrophils were stimulated *ex vivo* with PMA. We observed that HFD lung neutrophils cocultured with HFD LVECs secrete higher amount of ROS than those cocultured with CD LVECs (Fig. 5g). In a similar experimental setting, we cocultured CD or HFD LVECs with lung neutrophils isolated from CD mice, followed by *ex vivo* stimulation with PMA and quantification of ROS contents in the supernatants. We observed that CD lung neutrophils cocultured with HFD LVECs secrete higher amount of ROS than those cocultured with CD LVECs (Fig. 5h). These data thus suggest that HFD-exposed LVECs prime neutrophils for enhanced innate immune functions.

### LVECs activated by saturated fatty acids and cholesterol boost neutrophil functions

FRD has been related to increased serum fatty acids and cholesterol, which were shown in independent studies to activate vascular endothelial cells (VECs) ^35–41^. We therefore speculate that FRD-induced increased serum lipids stimulate LVECs to prime lung neutrophils. To evaluate the impact of FRD on serum lipid contents, we performed non-targeted metabolomics analysis in the sera of CD and HFD mice. We observed differences in serum metabolites, including 106 and 62 significantly altered metabolite contents in positive and negative ion modes, respectively (Extended Data Fig. 8a). Notably, 60% of these altered metabolites are lipids and lipid-like molecules (Extended Data Fig. 8b). Further lipidomics analysis showed that phosphatidylcholines (PC), triglyceride (TG), and sphingomyelin (SM) were among the most evidently changed lipid subtypes in the sera of HFD versus CD mice (Extended Data Fig. 8c). Although there were minimal changes in total contents of serum PC, lyso-phosphatidylcholine (LPC), TG, SM, and phosphatidylinositol (PI) in HFD versus CD mice (Extended Data Fig. 8d), there were evident changes in subtypes of lipids in all of these five lipid categories (Extended Data Fig. 8e-g).

In addition to HFD, we also observed evident changes in serum lipid contents in KD versus CD mice (Extended Data Fig. 8h). Similar to that in HFD mice, TG, PC, and SM were among the most evidently changed lipid subtypes in the sera of KD versus CD mice (Extended Data Fig. 8h), though KD was found to cause changes in more subtypes of serum lipids than HFD (Extended Data Fig. 8i-k). Further analysis on the contents of saturated versus unsaturated fatty acids revealed that both HFD and KD increased serum contents of saturated fatty acids (SFA), monounsaturated fatty acids (MUFA), and cholesterol (CL), while reducing polyunsaturated fatty acids (PUFA), as compared to those in CD mice (Fig. 6a-f). These data collectively suggest that FRD alters serum lipid contents characterized by increased levels of serum SFA/MUFA and CL, and reduced serum PUFA.

**Figure 6.**
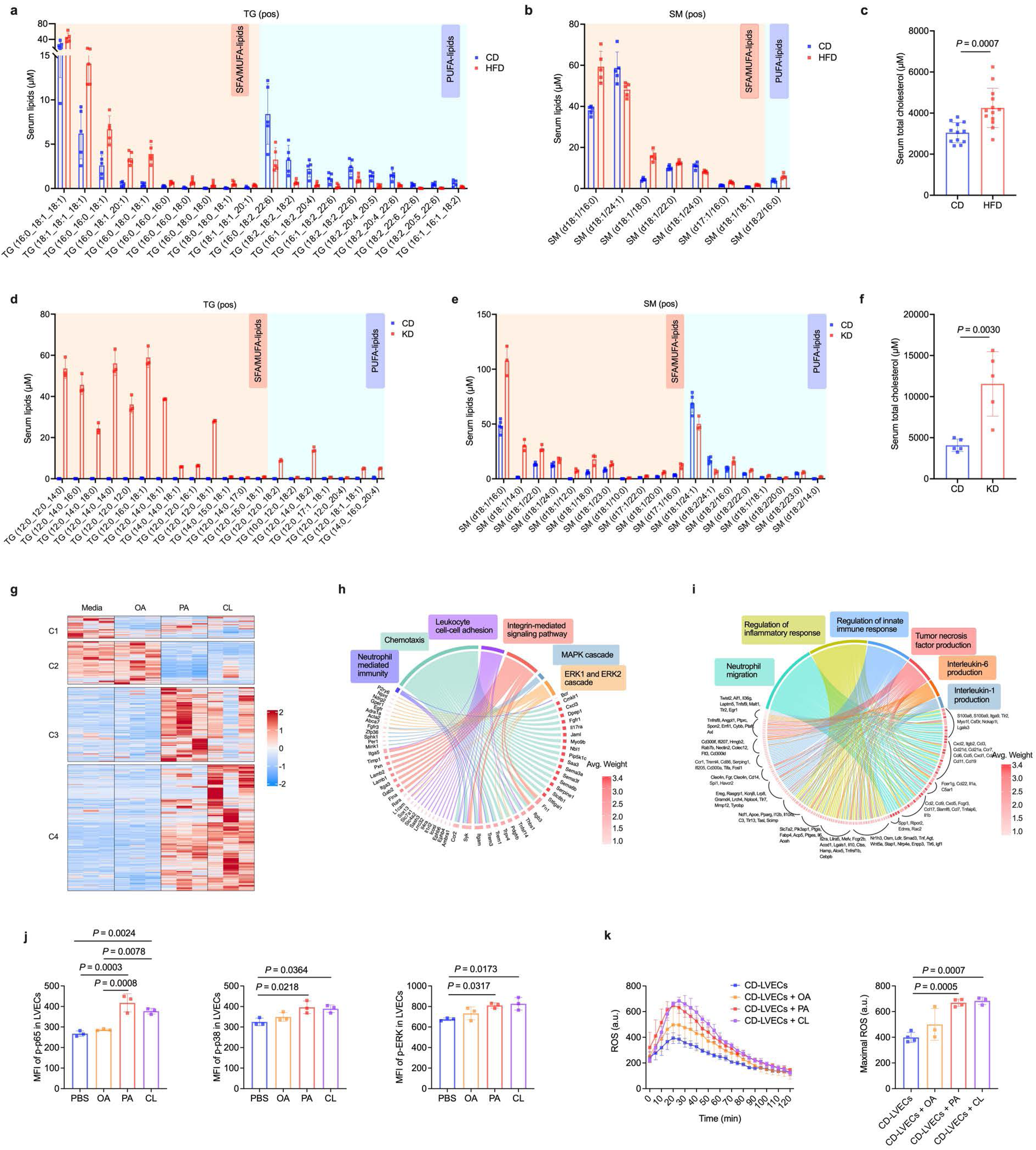
LVECs activated by saturated fatty acids and cholesterol boost neutrophil functions. (a) Quantification of the 20 most significantly altered TGs in serum of CD versus HFD mice, identified by positive-mode LC-MS. (b) Quantification of 8 representative SMs selected from the top 10 significantly altered SMs in serum between CD and HFD mice, detected via positive-mode LC-MS. (c) Total cholesterol contents in serum in CD and HFD mice. (d) Quantification of the 20 most significantly altered TGs in serum between CD and KD mice, identified by positive-mode LC-MS. (e) Quantification of the 19 significantly altered SMs in serum between CD and KD mice, detected via positive-mode LC-MS. (f) Total cholesterol contents in serum in CD and KD mice. (g) Heatmap showing differentially expressed genes (DEGs) in LVECs isolated from CD lung and treated with PA (0.1 mM), OA (0.1 mM), or CL (0.2 mM) for 3h compared to control cells without stimulation with lipids (media) (z-score normalized, P < 0.05). DEGs are grouped into four distinct clusters (C1-C4). (**h**,**i**) Chord diagram showing enriched GO pathways for DEGs in cluster 3 (**g**) and cluster 4 (**h**) related to cell adhesion, chemotaxis, immune response, and inflammatory signaling. Red colored bars indicate average statistical weight (-log10(p-value), 1.0-3.4) with deeper red showing higher significance. Colored lines connect genes to their associated pathways. (j) Median fluorescence intensity (MFI) of intracellular phosphorylated NF-κB p65 (p-p65), phosphorylated p38 MAPK (p-p38), and phosphorylated ERK (p-ERK) in LVECs isolated from CD lung. LVECs were treated with PA (0.1 mM), OA (0.1 mM), or CL (0.2 mM) for 3h before intracellular staining. (k) ROS contents in culture supernatants of PMA-stimulated CD lung neutrophils cocultured overnight with LVECs pretreated with palmitic acid (PA), oleic acid (OA), or cholesterol (CL). Left: kinetics of ROS contents in arbitrary units (a.u.). Right: peak ROS levels. n = 3 or 4 culture wells per group as indicated. Data in (**a**,**b**,**d**-**i**) are from one experiment with n = 3 or 5 mice per group as indicated. Data in (**c**) are pooled from two independent experiments (n = 12 in CD group and n = 13 in HFD group). Data in (**j**,**k**) are representatives of two independent experiments with n = 3 or 4 mice per group. Two-tailed Student’s t-test was used for comparison between two-groups. One-way ANOVA followed by a Tukey test was performed to compare more than two groups.

To directly address the potential stimulating effects of SFA, MUFA, and CL in LVECs, we cultured VEC cell lines including HBMEC and HUVEC, with palmitic acid (PA), Oleic acid (OA), or CL supplemented to the culture media. In both HBMEC and HUVEC, we observed time and dose-dependent increase of NF-κB p-p65, MAPK p-p38, and p-ERK after stimulation with PA, OA, and CL. Notably, PA and CL had overall more evident VEC-stimulating effects than OA (Extended Data Fig. 9a-d). To determine whether SFA, MUFA, and CL also activate primary mouse LVECs, we performed RNA sequencing analysis in LVECs isolated from CD mice and stimulated *ex vivo* with PA, OA, or CL. We observed that PA and CL induced evident transcriptional changes in mouse LVECs characterized by upregulated gene transcripts of pathways related to cell adhesion, chemotaxis, and inflammatory response, as compared to OA stimulation (Fig. 6g-i). PA and CL also showed more potent stimulating effects than OA in LVECs isolated from CD mice, as demonstrated by increased levels of intracellular NF-κB p-p65, MAPK p-p38, and p-ERK (Fig. 6j).

To further determine the roles of lipid-stimulated LVECs on neutrophils, we stimulated freshly isolated LVECs from CD mice with PA, OA, or CL. After removing lipid supplements from culture media, LVECs were cocultured overnight with freshly isolated lung neutrophils from CD mice, followed by PMA stimulation. We detected increased ROS contents in the culture supernatants of neutrophils cocultured with PA- and CL-stimulated LVECs, and to a less extent in those cocultured with OA-stimulated LVECs (Fig. 6k). These data collectively suggest that FRD-induced increased serum SFA and CL activate LVECs, which prime neutrophils for enhanced innate immune functions characterized by increased potential of ROS production.

### FRD-exposed LVECs boost neutrophil functions via contact- and ICAM-1-dependent mechanism

Lung neutrophils were shown to be in close contact with LVECs ^16, 17^. To determine whether FRD-exposed LVECs boost neutrophil functions via contact-dependent mechanisms, we utilized a trans-well system to separate lung neutrophils and LVECs in *ex vivo* coculture. We found that separation of neutrophils with LVECs abrogated the neutrophil-boosting effect by HFD LVECs, as evidenced by reduced ROS production by CD lung neutrophils cocultured with HFD LVECs (trans-well) to levels comparable to those cocultured with CD LVECs (Fig. 7a). These data suggest that FRD-exposed LVECs boost neutrophil functions via cell-cell contact-dependent mechanisms.

**Figure 7.**
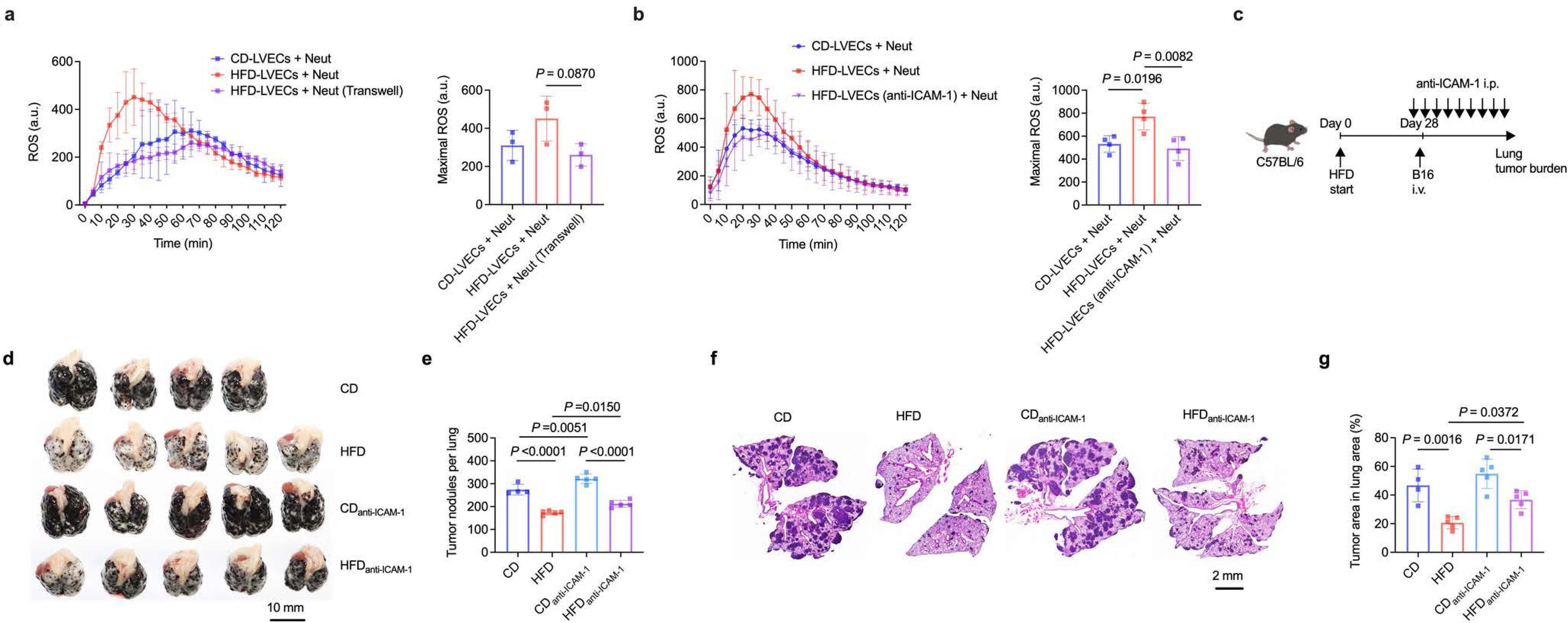
FRD-exposed LVECs boost neutrophil functions via contact- and ICAM-1-dependent mechanism. (a) ROS contents in culture supernatants of PMA-stimulated CD lung neutrophils cocultured overnight with CD or HFD LVECs prior to stimulation, with or without separation of neutrophils and LVECs by trans-well insertions. Left: kinetics of ROS contents in arbitrary units (a.u.). Right: peak ROS levels. n = 3 culture wells per group. (b) ROS contents in culture supernatants of PMA-stimulated CD lung neutrophils cocultured overnight with CD or HFD LVECs prior to stimulation, with or without pre-treatment of HFD LVECs with anti-ICAM-1 neutralizing antibody for 2 hours. Left: kinetics of ROS contents in arbitrary units (a.u.). Right: peak ROS levels. n = 4 culture wells per group. (c) Schema of the continuous *in vivo* ICAM-1 blockade in HFD and CD mice immediately before and after B16 melanoma cell inoculation i.v.. (**d**,**e**) Representative macroscopic lung images (**d**) and quantification of visible tumor nodules (**e**) on the surface of lung lobes. n = 4 or 5 mice per group as indicated. (**f**,**g**) Representative lung histopathology images (**f**) and percentage of lung area occupied by tumor lesions (**g**) based on lung histopathological analysis in mice shown in (**d**). Data are representative of two (**a**,**d**-**g**) or three (**b**) independent experiments. Graphs with error bars are presented as mean ± s.d. One-way ANOVA followed by a Tukey test was performed to compare more than two groups.

ICAM-1 play critical roles in the cell contact and interactions between neutrophils and VECs during various pathological scenarios ^42, 43^. To determine whether ICAM-1 is involved in the boosting effect of neutrophils by FRD-exposed LVECs, we incubated LVECs isolated from HFD mice *ex vivo* with anti-ICAM-1 blocking monoclonal antibodies (mAbs). After washing out the excessive mAbs, LVECs were cocultured with CD lung neutrophils, followed by PMA stimulation and quantification of ROS production. We found that blocking of ICAM-1 on LVECs abrogated the neutrophil-boosting effect by HFD LVECs (Fig. 7b), suggesting that FRD-exposed LVECs boost neutrophil functions via ICAM-1-dependent mechanisms. To further show the requirement of ICAM-1 in FRD-induced antitumor immunity in the lungs, we administered repeated doses of anti-ICAM-1 blocking mAbs *in vivo* to either CD or HFD mice immediately before and after inoculation i.v. with B16 melanoma cells (Fig. 7c). At experimental endpoint, we observed that *in vivo* blocking of ICAM-1 reduced HFD-induced pulmonary antitumor response (Fig. 7d-g). In contrast, blocking of ICAM-1 showed less evident effects in pulmonary antitumor response in CD mice, in particular when judged by the area of tumor lesions in lung histopathologic analysis (Fig. 7d-g). These data thus collectively suggest that FRD-exposed LVECs boost antitumor functions of lung neutrophils via ICAM-1-dependent mechanisms.

### Intensified cell adhesion and priming signals from human LVECs are associated with boosted functions in human intrapulmonary neutrophils

Our data thus far suggest that LVECs boosts lung neutrophil functions via intensified ICAM-1-dependent interactions in mice. We next sought to assess whether interactions between human intrapulmonary neutrophils and LVECs are associated with boosted defense functions in neutrophils. By using published original data on single-cell RNA sequencing (scRNA-seq) in human tissues including the lung, bone marrow, peripheral blood, and spleen^44^, we identified neutrophils from these tissues and found that human neutrophils showed tissue-specific transcriptional features (Fig. 8a,b), suggesting tissue adaptation of neutrophils which is consistent with observations in mouse neutrophils^9^. To determine the potential interactions between vascular endothelial cells and neutrophils in human lung tissues, we utilized publicly available datasets of large-scale scRNA-seq from non-small cell lung cancer (NSCLC) normal adjacent tissues^45, 46^, from which 6,370 neutrophils and 14,249 endothelial cells (ECs) were identified. Among these cells, we identified 7 clusters of neutrophils (N0-N6) and 12 clusters of ECs (clusters 0-11) (Fig. 8c,d). GO enrichment pathway analysis of differentially expressed genes in EC clusters showed that clusters 7 and 8 significantly upregulated transcription of genes related to chemotaxis, recruitment, and immune responses, indicating these ECs were in an activated state (Fig. 8e).

**Figure 8.**
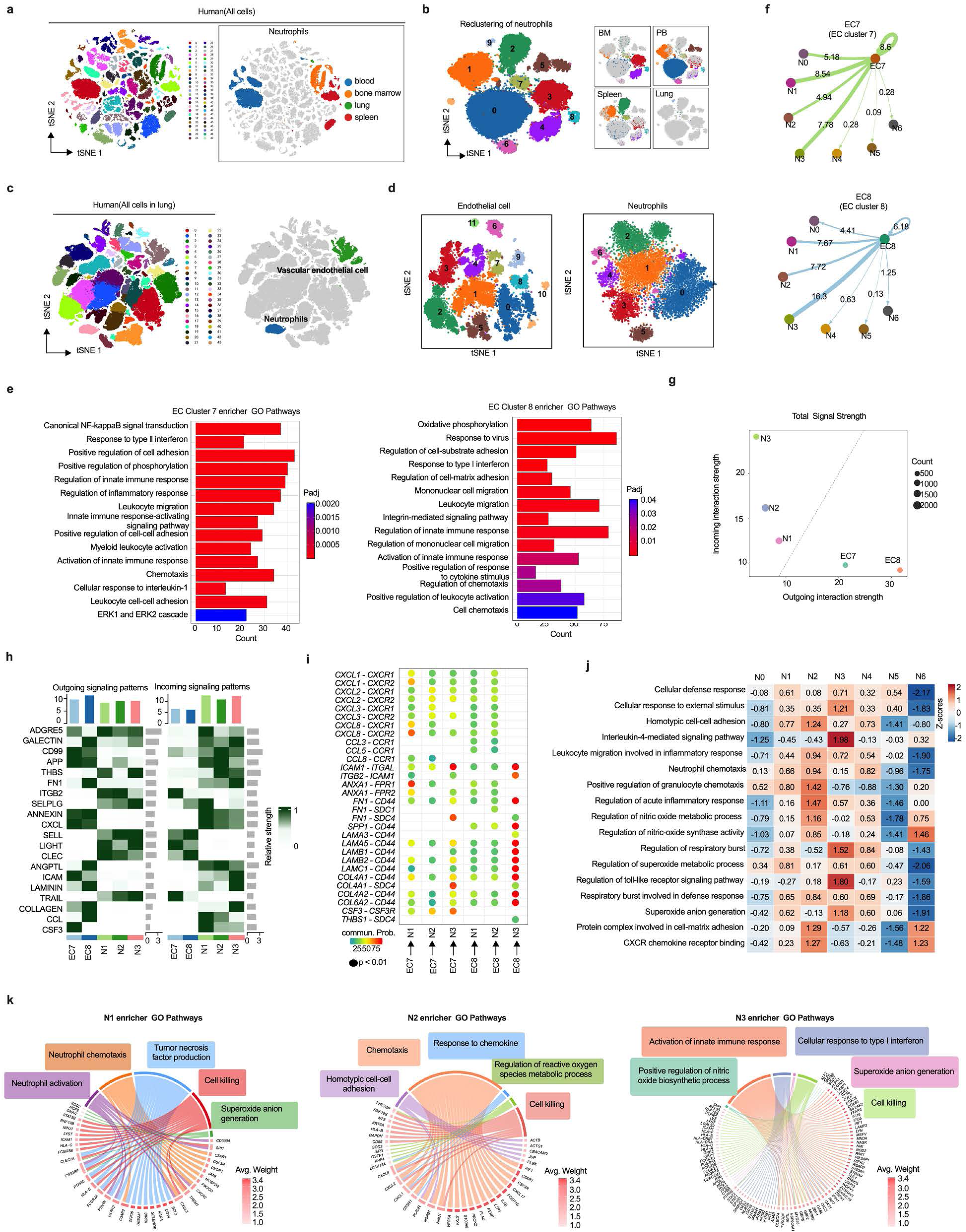
Intensified cell adhesion and priming signals from human LVECs are associated with boosted functions in human intrapulmonary neutrophils. (a) t-SNE plots depicting single-cell transcriptomes of human cells from bone marrow, blood, lung, and spleen (a total of 140,156 cells). Transcriptionally distinct clusters of neutrophils (right) from total cells (left) are highlighted. (b) t-SNE plots showing the transcriptionally distinct clusters of human neutrophils from various tissues. (c) t-SNE plots of single-cell transcriptomes from human lung cells (a total of 346,199 lung cells). Lung neutrophils and endothelial cells are highlighted (right). (d) t-SNE plots showing clusters of human lung endothelial cells (left) and neutrophils (right). (e) Gene ontology (GO) enrichment pathways of differentially expressed genes in endothelial cell clusters 7 and 8 as shown in (**d**). (f) Network plots visualizing total interaction strength between endothelial cell (EC) clusters 7 and 8 and clusters of neutrophils (N0 to N6) as shown in (**d**). Numbers on colored edges indicate quantified interaction strength between paired cell groups, with thicker lines indicating stronger interactions. (g) Scatterplot positioning dominant signaling sender (source) and receiver (target) cell populations based on interaction strength. Position of cell populations is determined by their outgoing (x-axis) and incoming (y-axis) interaction strengths, with circle size indicating cell count. (h) Heatmap illustrating signaling pathways contributing to outgoing (left) and incoming (right) signaling among specific cell groups. (i) Bubble plot visualizing significant ligand-receptor interactions from endothelial cell subpopulations (EC7 and EC8) to neutrophil subpopulations (N1, N2, and N3). (j) Heatmap displaying enrichment scores for 17 immune-related functional pathways across neutrophil subpopulations (N0 to N6). Color scale indicates standardized enrichment score. Numbers indicate the actual enrichment Z-score value. (k) Chord diagrams showing enriched GO pathways for differentially expressed genes in neutrophil subpopulations N1, N2, and N3 related to immune responses. Red bars indicate average statistical weight (-log10 of p-value).

Based on our observations in mice that FRD-activated LVECs boosts neutrophil antitumor functions via intensified ICAM-1-dependent cell-cell contact and signaling, we conducted an in-depth analysis on intercellular communication between human LVECs (clusters 7 and 8) and intrapulmonary neutrophils (clusters N0-N6). We conducted CellChat analysis and found that EC clusters 7 and 8 showed intensified interactions preferentially with neutrophils clusters N1-N3 (Fig. 8f). Analysis of outgoing and incoming signaling strengths showed that the two EC clusters primarily functioned as signal sources, while the three neutrophil clusters primarily served as signal receivers (Fig. 8g). Notably, we identified signaling pathways associated with cell-cell adhesion and chemotaxis, such as collagen family (COLLAGEN), C-X-C motif chemokine ligand family (CXCL), C-C motif chemokine ligand family (CCL), intercellular adhesion molecule family (ICAM), and annexin family (ANNEXIN), as key signals from EC clusters 7 and 8 to neutrophils N1-N3 (Figure 8h). Further examination of significant ligand-receptor pairs contributing to EC-to-neutrophil signaling identified 21 ligands in ECs and 11 receptors in neutrophils, including the interactions between EC-derived ICAM-1 and its receptors ITGAL on neutrophils (Fig. 8i), which is consistent with our data in mice demonstrating LVECs reprogram neutrophils via ICAM-1. Scoring analysis of activation and function-related pathways in neutrophil subtypes revealed that, among all pulmonary neutrophil clusters, clusters N1, N2, and N3 displayed enhanced immunological functions including chemotaxis and defense response (Figure 8j). GO enrichment pathway analysis of differentially expressed genes in N1, N2, and N3 neutrophils consistently showed enhancement in immune activation, killing capacity, and reactive oxygen species production (Figure 8k).

Collectively, these data suggest that human LVECs and neutrophils engage multiple ligand-receptor interactions, with chemokines, integrins, and adhesion molecules playing critical roles. Importantly, intensified LVECs-to-neutrophil adhesion and signaling are associated with functional reprogramming in neutrophils with enhanced antitumor innate immunity.

## Discussion

A recent milestone of research in neutrophils is their quick adaptation to tissue-specific signals in steady state in non-hematopoietic tissues including but not limited to the lung, skin, and intestine, which contributes to their functional diversification ^16^. Notably, neutrophils isolated from lung tissues of naïve mice demonstrate evident transcriptional, phenotypic, and functional differences against those in peripheral blood and the spleen ^16^. These findings underline the profound impact of lung tissue-specific microenvironment on neutrophils ^9^.

Updated evidence suggests that tissue adaptation of neutrophils in non-hematopoietic organs occurs primarily in the vasculature compartments of various organs ^1, 9^. In the lung, a non-negligible fraction of circulating neutrophils is entrapped in the vasculature, thus representing a marginal pool of neutrophils ^9, 47, 48^. A CXCR4 inhibitor, plerixafor, was shown to induce egress of marginal pool neutrophils from the lung to reenter the circulation ^47, 48^. A more recent study showed that CXCL12-expressing LVECs guide neutrophils to lung vascular niche for tissue-specific signals which is dependent on CXCR4 (receptor of CXCL12) on neutrophils ^16^. These findings collectively underscore the critical roles of LVECs in shaping neutrophil functions in steady-state lungs, though the significance of LVECs-induced tissue-specific reprograming of neutrophil functions in respiratory diseases, such as malignancies, has remained unclear. We show in this study that FRD activates LVECs, which subsequently reprograms the tissue adaptation of neutrophils with enhanced ROS production potential and tumor cytotoxic functions. Our findings therefore reveal previously unappreciated roles of lung tissue-specific signals in shaping neutrophil functions in respiratory diseases.

Previous studies suggest that neutrophils support lung colonization of metastatic breast cancer ^2, 3, 7, 8^. Obesity, either a genetic model (ob/ob mice) or induced in wild type mice with prolonged (up to 15 weeks) HFD, was shown to exacerbate metastasis of breast cancer cells to the lung in a neutrophil-dependent manner ^3, 8^. However, such a neutrophil-dependent prometastatic effect was independent on increased dietary fat contents but was dependent on increased fat body mass, as evidenced by increased lung metastasis in CD-fed ob/ob mice but not in HFD-fed Balb/c mice that are resistant to HFD-induced obesity ^8^. Moreover, obesity-induced systemic inflammation was responsible for such an exacerbated lung metastasis of breast cancer via disruption of vascular integrity and increased extravasation of tumor cells in the lung tissues ^3^. Importantly, obesity was shown to induce similar functional changes in both the lung and bone marrow neutrophils, suggesting a systemic rather than lung tissue-specific reprograming of neutrophils in obese mice ^3^. In contrast to these observations in obese mice, we show that FRD, including isocaloric HFD and KD, boost antitumor functions of lung neutrophils in a tissue-specific manner. The effect of FRD in our study was independent of excessive calorie intake or weight gain and was not associated with systemic reprogramming of neutrophils, nor was it dependent on disruption of endothelial integrity, as evidenced by comparable extravasation of tumor cells from the circulation to lung tissues in B16 i.v. model. This was further supported by FRD-induced anti-tumor response in an LLC lung cancer cell i.t. inoculation model which is independent of trans-endothelial extravasation of tumor cells. Instead, the effect of FRD was exerted via boosting antitumor potential of neutrophils in both prophylactic and therapeutic settings. Our findings therefore contrast those in previous studies in obese mice and thus shed light on a previously unappreciated mechanism through which dietary fat augments lung tissue-specific antitumor innate immunity which is independent on excessive calorie intake or gain of fat mass. Indeed, clinical observations suggest that in patients with non-small cell lung cancer, overweight but not obesity are related to favorable clinical outcomes ^49^. The roles of isocaloric diet with increased fat contents therefore deserve further investigation in cancer patients in particular in those with primary or metastatic cancers in the lungs ^50^.

To date, studies on VEC activation by lipids are mostly focused on its impact on atherosclerosis primarily involving arterial VECs, with the functional significance of its impact on VECs in the capillary much less addressed ^35–40, 51^. Notably, the interactions between neutrophils and LVECs were shown to be in the capillary of the lung vasculature ^16^. We show in this study that FRD-induced increased serum SFA/MUFA and cholesterol, which stimulate LVECs to have increased transcription of immune function-associated genes and activation of intracellular signaling pathways. Importantly, *ex vivo* stimulation of LVECs with SFA and cholesterol increased their neutrophil-stimulating functions. These findings shall foster future studies on the impact of serum lipids on VECs in the capillary, which likely contrasts that in atherosclerotic vascular lesions ^51^. Our study suggests that increased SFA and cholesterol favors the host via boosting the anti-tumor functions of lung neutrophils. In support of our findings, clinical evidence suggests that increased serum lipids, in particular cholesterol, are related to improved clinical outcomes in patients with lung cancer ^52, 53^.

Our findings highlight the functional significance of dietary fat-induced tissue-specific functional reprogramming of pulmonary neutrophils, which might be exploited to boost tissue-specific immunity against lung cancers. Our study shall foster future studies on diet-related tissue-specific reprograming of neutrophils in various diseases such as pulmonary infections, as well as in organs other than the lung, which will deepen our understanding on tissue-specific reprogramming of neutrophils and dietary intervention in patients with cancers and beyond.

## Acknowledgements

This work was supported by National Natural Science Foundation of China (32170893, 31970861, and 32470947 to Y.Y., 32100714 to Y.W., 32400742 to T.W., and 32171275 and 32371330 to Q.D.Z), Natural Science Foundation of Zhejiang Province (LR20H100001 and LRG25H100001 to Y.Y.), China Postdoctoral Science Foundation (2024M752824 to Y.W.), Postdoctoral Fellowship Program of CPSF (GZC20241486 to Y.W.), and Fundamental Research Funds for the Central Universities (226-2024-00161 to Q.D.Z.). We thank Z. Xing (McMaster University), Q. Wang and D. Wang (Zhejiang University) for proofreading the manuscript and helpful discussion. We thank X. Cao and Z. Guo (National Key Laboratory of Medical Immunology), Jianli Wang, Q. Wang, Feng Xu, X. Wang, Junxiao Wang, and Fei Xu (Zhejiang University), and J. Wang (Shanghai Jiao Tong University) for providing experimental materials or technical assistance. We thank Y. Li, J. Wang, and X. Yu from the Core Facilities of Zhejiang University School of Medicine for technical assistance.

## Author contributions

Y.W., Lei W., and Y.Y. conceived and designed the study. Y.W., J.Z., Y.L., T.W., Lu W., Y.M., C.C., Y.L., Z.H., Z.Y., H.X., and Q.D.Z. performed experiments. Y.W., Z.H., Z.Y., Q.D.Z., and Y.Y. analyzed the data. Y.W., Lei W., and Y.Y. wrote the paper.

## Materials and methods

### Animals and diets

Five-week-old female C57BL/6 mice were purchased from Slac Laboratory Animals. Ly6G-DTR mice on a C57BL/6 background were generated by Shanghai Model Organisms Center, and were generously provided by Dr. J. Wang (Shanghai Jiao Tong University). All animals were housed in a specific pathogen-free facility at Zhejiang University Laboratory Animal Center under controlled environmental conditions (12-hour light/dark cycle, 50-60% humidity, 20-25°C) with *ad libitum* access to standard chow and water. For all experiments, six-week-old female mice were randomly allocated to chow diet (CD, Xietong Shengwu) group, high-fat diet (HFD, Research Diets, Inc.) group, or ketogenic diet (KD) group. Two variants of KD were used: KD-1 containing 90% lard-derived fat and KD-2 containing 90% plant oil (custom-formulated by Xietong Shengwu). All animal procedures were performed in accordance with guidelines approved by the Animal Research and Ethics Boards of Zhejiang University.

### Cell lines

The GFP-expressing B16F10 cell line (B16-GFP) was generated from luciferase-expressing B16F10 cells (B16-luc; ATCC, CRL-6475-LUC2) by transduction with lentiviral vector encoding GFP under the EF-1α promoter, as previously described ^30^. B16F10, B16-luc, and B16-GFP cells were maintained in RPMI 1640 medium (Thermo Fisher Scientific) supplemented with 10% fetal bovine serum (FBS; Serana) and 1% penicillin/streptomycin (Biosharp). Lewis lung carcinoma cells (LLC; ATCC, CRL-1642) were cultured in DMEM medium (Thermo Fisher Scientific) supplemented with 10% FBS and 1% penicillin/streptomycin. Human umbilical vein endothelial cells (HUVEC; ATCC, PCS-100-010) and human cerebral microvascular endothelial cells (HBMEC; ATCC, CRL-3245) were maintained in Endothelial Cell Medium (ScienCell) supplemented with Endothelial Cell Growth Supplement, 5% FBS, and 1% penicillin/streptomycin. All cells were cultured at 37°C in a humidified atmosphere containing 5% CO_2_.

### Mouse tumor models

To establish experimental melanoma metastasis, B16 cells (5 × 10^5^) suspended in 200 μL phosphate-buffered saline (PBS) were administered via tail vein injection. For the orthotopic lung cancer model, LLC cells (5 × 10^5^) in 50 μL PBS were delivered to the lungs via intratracheal instillation. Tumor burden was assessed by measuring post-mortem lung weight and by histological analysis of lung sections using hematoxylin and eosin (H&E) staining. For the subcutaneous tumor model, B16 cells (5 × 10^5^) were inoculated subcutaneously into the abdominal flank. Subcutaneous tumor growth was monitored every 48 hours using a digital caliper, and tumor volume was calculated as V = (L × W^2^)/2, where L and W represent the longest and shortest diameters, respectively. Mice were euthanized at humane endpoints or between days 14 to 18 after tumor inoculation for tissue harvest.

### *In vivo* bioluminescence imaging

Bioluminescence imaging was performed using the IVIS Spectrum *in vivo* imaging system (PerkinElmer, Waltham, MA, USA). Mice were anesthetized via inhalation of 2% isoflurane in oxygen and maintained under anesthesia throughout the imaging procedure. D-Luciferin (150 mg/kg; Yeason Biotechnology) was administered intraperitoneally. Eight minutes post-injection, bioluminescent signals were acquired with a 40 second exposure time. White light and bioluminescent images were acquired and quantified using Living Image software (PerkinElmer).

### Lung macroscopic and histopathological analysis

Lung tissues were harvested at days 14 to 18 post tumor inoculation. Visible B16 tumor nodules on the surface of lung lobes were counted under a JSZ8 stereomicroscope (Jiangnan Novel Optics, Nanjing, China). The lung tissues were then fixed in 4% paraformaldehyde for 48 hours at 4°C, dehydrated through graded ethanol series, embedded in paraffin, and sectioned at 5 μm thickness. H&E-stained lung sections were digitally scanned using a VS200 slide scanner (Olympus, Tokyo, Japan) and analyzed by using OlyVIA 4.1 software (Olympus). For confocal microscopy analysis, paraffin-embedded lung tissue sections underwent deparaffinization and rehydration before immunohistochemical staining. Heat induced antigen retrieval was performed using citrate antigen retrieval solution (pH 6.0; Solarbio). The sections were then permeabilized with 0.5% Triton X-100 in PBS for 10 minutes at room temperature. Following permeabilization, the sections were blocked with a solution containing 5% bovine serum albumin and 5% normal goat serum in PBS for 1 hour at room temperature. The blocked sections were sequentially incubated with primary antibodies overnight at 4°C, followed by secondary antibodies for 1 hour at room temperature. Specimens were mounted using DAPI Fluoromount-G (SouthernBiotech) according to the manufacturer’s protocol. For staining of Ly6G, anti-mouse Ly6G (clone 1A8; BioXCell) was used as the primary antibody, followed by goat anti-rat AF633 (Thermo Fisher) as the secondary antibody. Images were acquired using a Zeiss LSM 880 confocal microscope equipped with ZEN Black imaging software (Zeiss).

### *In vivo* cell depletion and neutralization of ICAM-1

For CD4 and CD8 T cell depletion *in vivo*, mice were injected intraperitoneally (i.p.) with anti-CD4 (clone GK1.5, BioXCell) and anti-CD8 (clone 2.43, BioXCell) at 200 μg each per animal. NK cells were depleted by i.p. administration of anti-NK1.1 (clone PK136, produced in-house, 200 μg per animal). For continuous T cell or NK cell depletion, repeated doses of 100 μg of depleting antibodies were administered at 7-day intervals through the experimental endpoint. Alveolar macrophages (AMs) were depleted using a single dose of clodronate liposomes or control liposomes (Liposoma Technology, 100 μL per mouse) administered intratracheally (i.t.) in two split doses. For circulating monocyte depletion, mice receiving i.t. clodronate/control liposomes were administered intravenously (i.v.) with clodronate or control liposomes (100 μL per mouse) at 5-day intervals, starting from 2 days post-initial i.t. injection through experimental endpoint. Neutrophil depletion was achieved by i.p. administration of anti-Ly6G monoclonal antibody (clone 1A8, BioXCell, 200 μg per animal), followed by repeated doses of 100 μg every 3 days. In Ly6G-DTR mice, diphtheria toxin (DT; ListLabs) was administered i.p. at 20 μg/kg every 2 days. For *in vivo* blocking of ICAM-1, 100 μg of anti-ICAM-1 antibody (clone YN1/1.7.4, BioXCell) was administered i.p. daily through endpoint.

### Isolation of immune cells and lung vascular endothelial cells

Cells from peripheral blood, spleen and lung tissues were isolated as previously described ^30^. Briefly, peripheral blood (500 to 600 μL) was collected from the abdominal vein and diluted with 2 mL PBS containing 15 mM EDTA (Thermo Fisher Scientific). Following centrifugation at 500 × g for 5 minutes at 4°C, cell pellets underwent erythrocyte lysis by resuspension in 3 mL RBC lysis buffer (SolarBio) for 3 minutes at room temperature. The reaction was terminated by adding 10 mL RPMI medium supplemented with 2% FBS, followed by centrifugation at 500 × g for 5 minutes at 4°C. After a second round of erythrocyte lysis, cells were resuspended in PBS. For peripheral blood neutrophil isolation, cells were labeled with anti-CD45 (APC-Cy7, clone 30-F11, BD Biosciences), anti-CD11b (PE-Cy7, clone M1/70, BD Biosciences), and anti-Ly6G (BV650, clone 1A8, BioLegend), followed by flow cytometry sorting of CD45^+^ CD11b^+^ Ly6G^+^ neutrophils on a BD FACS Aria Fusion cell sorter (BD Biosciences).

For spleen cell isolation, spleens were mechanically dissociated through a 70 μm cell strainer (BD Biosciences) in 3 mL PBS using a syringe plunger. The resulting single-cell suspension was centrifuged at 500 × g for 5 minutes at 4°C. Cell pellets were subjected to erythrocyte lysis using 2 mL of lysis buffer for 2 minutes at room temperature, washed with PBS, and resuspended in 500 μL of PBS. To isolate splenic neutrophils, a two-step sorting strategy was exploited. Splenic cells were stained with anti-CD3 Biotin antibody (clone 145-2C11, BioLegend) and anti-B220 Biotin antibody (clone RA3-6B2, BioLegend), followed by labeling with anti-Biotin microbeads (Miltenyi Biotec) for negative magnetic selection using a MS magnetic purification system (Miltenyi Biotec). CD3^-^B220^-^ splenic cells were labeled with anti-CD45, anti-CD11b and anti-Ly6G antibodies for flow cytometry cell sorting of neutrophils as described above.

For lung cell isolation, lung lobes were harvested, cut into small pieces, and enzymatically digested in collagenase type I (Thermo Fisher) for 1 hour at 37°C in a shaker incubator at 200 rpm. The digested lung tissues were crushed and filtered through a 100 μm cell strainer (BD Biosciences) to obtain a single-cell suspension. Following erythrocyte lysis, neutrophils were isolated either by flow cytometric sorting of CD45^+^ CD11b^+^ Ly6G^+^ cells or by positive selection using Ly6G microbeads (Miltenyi Biotec) on a LS magnetic purification system (Miltenyi Biotec). To isolate lung endothelial cells (LVECs), lung lobes were enzymatically digested in dispase (Sigma-Aldrich) for 40 minutes at 37°C. CD45^-^ CD31^+^ LVECs were sorted on BD FACS Aria Fusion cell sorter (BD Biosciences). To obtain purified AMs, bronchoalveolar lavage (BAL) cells were labeled with anti-CD11c microbeads (Miltenyi Biotec) followed by positive selection on a MS magnetic purification system (Miltenyi Biotec) according to the manufacturer’s instructions.

### Flow cytometry

Flow cytometry analyses were performed as previously described ^30^. Briefly, single-cell suspensions obtained from peripheral blood, the spleen, and the lung were seeded in U-bottom 96-well cell culture plates at ≤ 2 × 10^6^ cells per well in PBS. Cell viability was assessed using the Zombie Aqua fixable viability kit (BioLegend) according to the manufacturer’s instructions. Cells were then washed with PBS and incubated with anti-CD16/CD32 (clone 2.4G2, BD Biosciences, 1:200) in PBS containing 0.5% bovine serum albumin for 15 minutes on ice. Cells were labeled with the following fluorochrome-conjugated antibodies at 4°C for 30 minutes: anti-CD45 APC-Cy7 (clone 30-F11, BD Biosciences, 1:400), anti-CD45.2 PerCP-Cy5.5 (clone 104, Biolegend, 1:200), anti-CD45 BUV395 (clone 30-F11, eBioscience, 1:200), anti-CD11b PE-Cy7 (clone M1/70, BD Biosciences, 1:400), anti-CD11b BV605 (clone M1/70, Biolegend, 1:500), anti-Ly6C BV711 (clone HK1.4, Biolegend, 1:500), anti-Ly6G BV650 (clone 1A8, Biolegend, 1:500), anti-Siglec-F PE-CF594 (clone E50-2440, BD Biosciences, 1:500), anti-CD177 AF647 (clone Y127, BD Biosciences, 1:200), anti-CD14 APC-Cy7 (clone Sa14-2, Biolegend, 1:200), anti-CD49d FITC (clone R1-2, Biolegend, 1:200), anti-CD54 PE (clone YN1/1.7.4, Biolegend, 1:200), anti-CX3CR1 BV785 (clone SA011F11, Biolegend, 1:200), anti-CD31 PE (clone 390, Biolegend, 1:400), and anti-CD31 Pacific Blue (clone 390, Biolegend, 1:400).

For phospho-flow cytometry analysis, purified LVECs (2 to 5 × 10^5^ cells) were seeded in U-bottom 96-well cell culture plates and were stimulated *ex vivo* with palmitic acid (PA, 0.1 mM; Sigma-Aldrich), oleic acid (OA, 0.1 mM; Sigma-Aldrich), or cholesterol (0.2 mM; Beyotime) for 3 hours at 37°C. Cells were then fixed and permeabilized using fixation/permeabilization buffer (Thermo Fisher Scientific) according to the manufacturer’s instructions, followed by staining with the following antibodies for 30 min at room temperature: anti-ERK1/2 Phospho (Thr202/Tyr204) BV421 (clone 6B8B69, BioLegend, 1:10), anti-p38 MAPK Phospho (Thr180/Tyr182) PE (clone A16016A, BioLegend, 1:10), or anti-NF-κB p65 Phospho (Ser536) (93H1, Cell Signaling Technology, 1:1000) followed by Alexa Fluor 647-conjugated anti-rabbit secondary antibody (Thermo Fisher Scientific, 1:500).

For intravascular staining, 2 μg of anti-CD45 APC-Cy7 (clone 30-F11, BD Biosciences) diluted in 100 μL of PBS was injected i.v. via the tail vein of anesthetized mice. Organs were harvested 3 minutes post-injection.

Flow cytometric analysis was performed using a BD LSRFortessa flow cytometer equipped with FACS Diva software (version 8.0.1, BD Biosciences). Data analysis and visualization were conducted using FlowJo software (version 10.7.1, BD Biosciences).

### RNA-seq and data analysis

Neutrophils isolated from peripheral blood, spleen, and lung tissues, and isolated LVECs were pelleted and lysed in TRIzol reagent (Thermo Fisher Scientific), followed by storage at - 80°C until RNA extraction. Quality and quantity of extracted RNA were assessed using an N50 NanoPhotometer microvolume spectrophotometer (IMPLEN) and an RNA Nano 6000 assay kit on the Bioanalyzer 2100 system (Agilent Technologies). RNA sequencing libraries were prepared using the NEBNext Ultra RNA Library Prep Kit (New England Biolabs) following the manufacturer’s protocol, and unique index codes were added to identify individual samples. The index-coded samples were clustered using a TruSeq PE Cluster Kit v3-cBot-HS (Illumina) on a cBot cluster generation system according to the manufacturer’s guidelines. Following cluster generation, the library preparations were sequenced on an Illumina NovaSeq platform.

Total RNA-seq data in FASTQ format were preprocessed using custom Perl scripts to obtain clean reads by eliminating adapter sequences, poly(N) reads, and low-quality reads. The filtered reads were aligned to the mouse reference genome (mm10) obtained from the University of California Santa Cruz (UCSC) Table Browser using HISAT2 (version 2.0.5). Read counts for each gene were quantified using featureCounts (version 1.5.0-p3). Gene expression levels were normalized and calculated as Fragments Per Kilobase of transcript per Million mapped reads (FPKM) based on the gene length and mapped read counts. Data analysis was performed using R (version 4.3.3) and Python (version 3.9.7). Principal Component Analysis (PCA) was performed using the prcomp function in R with scaled and centered data. Differential expression analysis was conducted using the DESeq2 package (version 1.46.0) in R. P values were adjusted for multiple testing using the Benjamini-Hochberg procedure to control the false discovery rate (FDR). Genes with an adjusted P value < 0.05 were considered significantly differentially expressed.

For visualization of differential expression analysis, volcano plots were constructed using matplotlib (version 3.8.0) in Python, with data preprocessing performed using pandas (version 2.1.4) and numpy (version 1.26.4) packages. Hierarchical clustering and heatmap visualization were performed using the ComplexHeatmap package (version 2.18.0) in R, with color scaling and additional graphical elements implemented through the circlize (version 0.4.16) and grid (version 4.3.3) packages. The overlapping patterns of differentially expressed genes (DEGs) were illustrated using proportional Venn diagrams generated by the matplotlib-venn package (version 0.11.7). Gene Ontology (GO) enrichment analysis was conducted using the ClusterProfiler package (version 4.2.1) with the org.Mm.eg.db annotation database (version 3.17.0). GO terms were considered significantly enriched at an adjusted P-value threshold of 0.05.

### Enzyme-linked immunosorbent assay (ELISA)

Sorted neutrophils were stimulated *ex vivo* with phorbol-12-myristate-13-acetate (PMA, Aladdin) at a concentration of 50 or 100 ng/mL, or LPS at 5 or 10 ng/mL. After incubation for 12 hours, cell culture supernatants were harvested for cytokine analysis. The levels of tumor necrosis factor (TNF), interleukin-1β (IL-1β), and interleukin-6 (IL-6) were determined using Mouse ABTS ELISA development kits (Peprotech) according to the manufacturer’s instructions.

Absorbance was measured at 405 nm with wavelength correction at 650 nm on a Synergy Mx M5 microplate reader (Molecular Devices). Cytokine concentrations were calculated using standard curves generated from serial dilutions of recombinant proteins using GraphPad Prism software (version 9.0, GraphPad Software).

### Assay of tumor cytotoxicity

Splenic and lung neutrophils or AMs, isolated as described above, were cocultured with B16-luc melanoma cells in complete culture medium for 48 hours. Culture supernatants were collected for cytokine analysis using ELISA. Viable B16-luc tumor cells were quantified by measuring luciferase activity in cell lysates using a luciferase reporter assay system (Yeason Biotechnology) on a Synergy Mx M5 plate reader (Molecular Devices) following the manufacturer’s instructions. For reactive oxygen species (ROS) inhibition studies, selected cell culture wells were supplemented with 5 mM N-acetyl-cysteine (NAC; Sigma-Aldrich).

### Detection of ROS

Isolated neutrophils were resuspended to 5 × 10 cells in 100 μL serum-free media (DMEM-F12 without phenol red; Gibco) and plated in white flat-bottom 96-well plates (Thermo Fisher Scientific). Luminol (Sigma-Aldrich) was added to a final concentration of 100 μM. Cells were stimulated either by PMA (Aladdin) at 100 nM or by coculture with B16-luc cells at a 5:1 neutrophil-to-tumor cell ratio. Prior to measurement of ROS, plates were gently agitated, and luminescence was measured across all wavelengths using a Synergy Mx M5 microplate reader (Molecular Devices) at 5-minute intervals for 100 or 120 minutes. For comparative analyses between splenic and lung neutrophils, relative ROS production was calculated by subtracting baseline luminescence values (unstimulated cells) from those of their corresponding stimulated cells.

For coculture studies, neutrophils (1 × 10 cells/well) were incubated overnight with LVECs at a 1:2 ratio in a final volume of 200 μL. After washing, luminol was added, and cells were stimulated with 100 nM PMA. In trans-well experiments, neutrophils and LVECs were separated using 0.4 μm pore size trans-well inserts (Corning) in 24-well tissue culture plates.

For lipid stimulation experiments, LVECs were treated *ex vivo* for 3 hours with either palmitic acid (PA; 0.1 mM; Sigma-Aldrich), oleic acid (OA; 0.1 mM; Sigma-Aldrich), or cholesterol (0.2 mM; Beyotime). After removing lipids from culture media, LVECs were cocultured with neutrophils overnight. For ICAM-1 blocking experiments, LVECs were pre-incubated with anti-ICAM-1 antibody (clone YN1/1.7.4; BioXCell; 5 μg/mL) for 2 hours, washed to remove excessive ICAM-1 in culture media, and cocultured overnight with neutrophils. Cells were then stimulated with 100 nM PMA for ROS production analysis.

### LC-MS/MS analysis of serum metabolome

Serum was obtained by collecting blood samples from mice and allowing them to clot for 60 minutes at room temperature. Samples were centrifuged at 3,000 × g for 15 minutes at 4°C, and the resulting serum was stored at -80°C until analysis. Metabolite extraction was performed by mixing 100 μL of serum with 400 μL of 80% LC-MS grade methanol. The mixture was vigorously vortexed for 30 seconds and incubated on ice for 5 minutes, followed by centrifugation (15,000 × g, 4°C, 20 minutes). The supernatant was collected and diluted with LC-MS grade water to a final methanol concentration of 53%. Quality control (QC) samples were prepared by pooling equal volumes of all serum samples. Metabolomic analysis was performed using a Vanquish UHPLC system (Thermo Fisher Scientific) coupled to a Q Exactive HF-X mass spectrometer. Chromatographic separation was achieved using a Hypersil Gold C18 column (100 × 2.1 mm, 1.9 μm) maintained at 40°C. The mobile phase flow rate was set at 0.2 mL/min. For positive ionization mode, mobile phase A consisted of 0.1% formic acid in water, while mobile phase B was methanol. For negative ionization mode, mobile phase A was 5 mM ammonium acetate (pH 9.0), and mobile phase B was methanol. Mass spectrometry was performed in both positive and negative electrospray ionization (ESI) modes.

Raw data files were processed using CD 3.1 software with the following parameters: retention time tolerance of 0.2 min, mass accuracy of 5 ppm, signal intensity variation of 30%, and signal-to-noise ratio of 3. Metabolite identification was conducted by comparing accurate mass measurements and MS/MS fragmentation patterns against mzCloud, mzVault, and Masslist databases. Peak areas were normalized using QC samples, and features exhibiting a coefficient of variation (CV) greater than 30% in QC samples were excluded from further analysis.

Metabolites were annotated using the KEGG, HMDB, and LIPIDMaps databases. Differential metabolites were visualized using volcano plots with log2 fold change on the x-axis and -log10(P-value) on the y-axis. Metabolites were considered significantly changed when meeting both |log2 fold change| > 0.58 and P value < 0.05 (Student’s t-test, adjusted for multiple comparisons using Benjamini-Hochberg method). The point size in the plot was scaled according to the Variable Importance in Projection (VIP) score from the OPLS-DA model. Volcano plots were generated using Python (version 3.9.7) with matplotlib package.

### Serum lipid extraction and lipidomics analysis

Serum lipids were extracted using a modified methyl tert-butyl ether (MTBE)-based protocol as previously described ^54^. Briefly, aliquots of mouse serum (20 μL) were transferred into plasticizer-free Eppendorf Safe-Lock microcentrifuge tubes (1.5 mL, Sigma-Aldrich). Each sample was supplemented with 225 μL of ice-cold methanol containing SPLASH LipidoMIX Internal Standard mixture, followed by addition of 750 μL MTBE. The mixtures were thoroughly vortexed for 30 seconds and incubated at room temperature (25-30°C) for 30 minutes to ensure complete lipid extraction. Phase separation was initiated by adding 188 μL of ultrapure water (Milli-Q) to each sample, followed by vigorous vortexing for 30 seconds. The samples were then centrifuged (13,000 × g, 15 minutes) to achieve distinct phase separation. The upper organic phase was carefully collected, transferred to fresh microcentrifuge tubes, and concentrated to dryness using a vacuum centrifuge. The dried extracts were stored at -80°C until analysis.

For LC-MS/MS analysis, dried lipid extracts were reconstituted in 200 μL of dichloromethane: methanol (1:1, v/v) containing 10 mM ammonium acetate. After centrifugation (13,000 × g, 15 minutes), the supernatants were transferred to 2 mL borosilicate glass vials. Lipidomic analysis was performed using a Shimadzu UPLC system coupled to a ZenoTOF 7600 mass spectrometer (SCIEX). Lipid separation was achieved on a Phenomenex Kinetex C18 column (100 × 2.1 mm, 2.6 μm) using mobile phase A (methanol: acetonitrile:water, 1:1:1, v/v/v, containing 5 mM ammonium acetate) and mobile phase B (isopropanol containing 5 mM ammonium acetate). Mass spectrometric data were acquired using information-dependent acquisition (IDA) in both positive and negative ionization modes.

Data processing and lipid species identification were performed using MS Dial 5.1. Individual lipid species were quantified by comparing their peak areas to those of corresponding SPLASH LipidoMIX Internal Standards [15:0-18:1(d7) PC, 15:0-18:1(d7) PE, 15:0-18:1(d7) PS, 15:0-18:1(d7) PG, 15:0-18:1(d7) PI, 15:0-18:1(d7) PA, 18:1(d7) LPC, 18:1(d7) LPE, 18:1(d7) Chol Ester, 15:0-18:1(d7) DAG, 18:1(d7) MAG, 15:0-18:1(d7) TAG, d18:1-18:1(d9) SM and Cholesterol (d7)]. All measurements were normalized to serum volume.

Lipid expression profiles were analyzed and visualized using R (version 4.3.3) and Python (version 3.9.7). To evaluate metabolic differences between CD and KD or HFD groups, principal component analysis (PCA) was performed using the ggplot2 and dplyr packages in R. Hierarchical clustering analysis and heatmap visualization were conducted using the ComplexHeatmap package in R. Differential lipid analysis was visualized using volcano plots, with significance thresholds set at |log2(HFD/CD) | or |log2(KD/CD) | > 0.5 and p < 0.05. The distribution of double bonds across different lipid classes was analyzed using Python. Within each lipid class, the relative abundance was calculated by normalizing the intensity of individual lipid species to the total intensity of that class. These distributions were visualized as stacked bar plots representing the fractional distribution of double bonds.

### Quantification of serum total cholesterol

Total cholesterol levels in mouse serum were measured using the Amplex Red Cholesterol and Cholesteryl Ester Assay Kit (Beyotime) according to the manufacturer’s instructions. Briefly, serum samples were diluted 100-fold with assay buffer, and 50 μL of diluted sample were mixed with 50 μL of working solution containing Amplex Red, cholesterol esterase, and enzyme mix. The reaction mixture was incubated at 37°C for 30 minutes in the dark. The absorbance was measured at 570 nm using a Synergy Mx M5 microplate reader (Molecular Devices).

### Immunoblotting

HUVEC or HBMEC cells were treated with PA, OA, or cholesterol for specified time periods and at various concentrations as indicated in figures. Total protein was extracted using RIPA lysis buffer (Beyotime) supplemented with phenylmethanesulfonyl fluoride (PMSF) (Beyotime) and phosphatase inhibitors (Beyotime). Proteins were separated by SDS-PAGE and transferred to PVDF membranes. The membranes were blocked with 5% non-fat milk and then incubated with primary antibodies overnight at 4°C, followed by incubation with corresponding secondary antibodies for 1 hour at room temperature. After three 10-minute washes, the membranes were incubated with HRP (Horseradish Peroxidase) chemiluminescent substrate (Fdbio Science) and imaged using a Tanon 4500 Gel Imaging System. For reprobing, membranes were stripped using stripping buffer for 20 minutes at room temperature, washed three times with TBST, and then reblocked with 5% non-fat milk before incubation with another primary antibody. The following primary antibodies were used: anti-NF-κB p65, anti-phospho-NF-κB p65, anti-p38 MAPK, anti-phospho-p38 MAPK, anti-β-actin (Cell Signaling Technology), anti-phospho-ERK1/2, anti-tubulin (Abmart), and anti-ERK1/2 (Proteintech).

### Real-time quantitative PCR with reverse transcription (RT-qPCR)

Total RNA was extracted from lungs using TRIzol reagent (Thermo Fisher Scientific). Reverse transcription was performed using PrimeScript RT Master Mix (TaKaRa Bio) following the manufacturer’s instructions. Quantification of B16-luc melanoma cells in the lungs was performed by RT-qPCR analysis of genes encoding premelanosome protein (*Pmel*), dopachrome tautomerase (*Dct*), and luciferase. Gene expression levels were normalized to *Gapdh* using the 2^-ΔΔCt method. The following primers (Tsingke Biotechnology) were used for amplification: *Pmel* (forward: 5′-GCTTGTAGGTATCTTGCTGGTGTT-3′, reverse: 5′- CCTGCTTCTTAAGTCTATGCCTATG-3′), *Dct* (forward: 5′-GGCTACAATTACGCCGTTG- 3′, reverse: 5′-CACTGAGAGAGTTGTGGACCAA-3′), luciferase (forward: 5′- CACCGTCGTATTCGTGAGCA-3′, reverse: 5′-AGTCGTACTCGTTGAAGCCG-3′), and *Gapdh* (forward: 5′-GTCGTGGAGTCTACTGGTGTCTT-3′, reverse: 5′- GTCATATTTCTCGTGGTTCACACCC-3′).

### Analysis of human scRNA-seq datasets

Single-cell RNA sequencing (scRNA-seq) data from human donor-derived bone marrow (BM), peripheral blood (PB), lung, and spleen tissues were obtained from three donors (TSP2, TSP14, and TSP25) through the Tabula Sapiens consortium data portal (https://tabula-sapiens-portal.ds.czbiohub.org/). Human lung scRNA-seq data were acquired from the Zenodo repository (https://doi.org/10.5281/zenodo.6411867), with a subset of 139 samples from normal lung tissues adjacent to tumor sites selected for analysis.

Quality control and data preprocessing were performed using Scanpy (v1.11.0). We implemented stringent filtering criteria, retaining cells with greater than 200 unique molecular identifiers (UMIs) and less than 20% mitochondrial gene expression. Count data were normalized to a total sum of 10,000 counts per cell using Scanpy’s normalize_total function with parameter target_sum=1e4, followed by natural logarithm transformation (log1p). Highly variable genes were identified using the Seurat v3 methodology, selecting the top 3,000 genes for downstream analysis. Technical covariates, including total UMI count and mitochondrial percentage, were regressed out, and the resulting matrix was scaled using Scanpy’s scale function with parameter max_value=10 to prevent outlier dominance. Principal component analysis (PCA) was subsequently performed, retaining the top 50 principal components. To mitigate batch effects, Harmony (v0.0.10) was applied to the PCA embeddings, generating batch-corrected representations of the data. Subsequently, a k-nearest neighbor graph was constructed from the corrected components, followed by t-SNE visualization and Leiden clustering analysis.

Cell clusters were annotated using SingleR (v2.8.0) with the BlueprintEncodeData() reference dataset, then cross validated against the cell type assignments from original publications. We next identified cluster specific marker genes with Scanpy’s tl.rank_genes_groups() function and compared them to known markers from the CellMarker database (http://bio-bigdata.hrbmu.edu.cn/CellMarker/). Neutrophil clusters were defined by high *FCGR3B (CD16b)*, *CXCR2*, *CSF3R*, and *ARG1* expression, whereas endothelial clusters expressed *VWF*, *CLDN5*, and *CDH5*.

### Differential gene expression analysis in human lung scRNA-seq data

Differential gene expression analysis was conducted using the Wilcoxon rank-sum test as implemented in Scanpy’s rank_genes_groups function. To account for multiple hypothesis testing, p-values were adjusted using the Benjamini-Hochberg procedure to control the false discovery rate (FDR). Genes with adjusted p-values < 0.05 were designated as differentially expressed genes (DEGs). For downstream analysis, upregulated DEGs (adjusted p-value < 0.05, log2 fold change > 0) were subjected to Gene Ontology (GO) enrichment analysis using the clusterProfiler package (v4.14.6) in R.

### Endothelial-neutrophil cell communication analysis

We used a multi-level computational framework to analyze crosstalk between lung endothelial cells and neutrophils. First, we systematically cataloged potential ligand-receptor interaction pairs between these cell populations using the human CellChatDB database. To integrate heterogeneous datasets, we implemented a deep learning-based variational autoencoder (scVI with n_latent=30, 1-3 hidden layers, trained for max_epochs=400) with batch correction, creating a unified latent representation that preserves cell-specific molecular signatures while enabling cross-cell type comparisons. Within this latent embedding, we identified signaling-enriched cells by selecting those with ligand or receptor expression above the 90th percentile, then utilized a fine-grained nearest-neighbor search algorithm (k=5) to identify potential communicating cell pairs with complementary signaling molecule expression. The communication scoring system integrated multiple parameters: molecular expression levels (base score calculated as ligand×receptor expression), cellular proximity (similarity score derived as exp(-0.5×distance) in latent space), and network context (reflecting local connectivity patterns). These components were weighted and combined into a composite confidence metric that was further refined through graph-based diffusion analysis (diffusion coefficient=0.5). Finally, we constructed a directed communication network between endothelial cells and neutrophils, with edge weights representing both molecular interaction strength and cellular spatial relationships, identifying predominant signaling pathways between these cell populations.

To visualize communication between endothelial and neutrophil clusters, we rendered network diagrams in R with igraph (v2.1.4), scatterplots and ligand-receptor dot plots with ggplot2 (v3.5.2), and ligand-receptor pathway activity heatmaps with ComplexHeatmap (v2.22.0) and circlize (v0.4.16).

### Neutrophil pathway activity assessment

Gene sets relevant to neutrophil biology were curated from the GO Biological Process collection (C5) accessed via the msigdbr R package (v10.0.2). To ensure statistical robustness, gene sets were filtered to include only those containing a minimum of five genes detected within our single-cell RNA sequencing dataset. Pathway activity was quantified using the AddModuleScore function implemented in the Seurat package to compute module scores for each pathway in individual cells, which were then aggregated at the cluster level. The resulting cluster-by-pathway matrix was row-wise Z-score normalized and visualized as a heatmap using pheatmap (v1.0.12) with a divergent color scale to highlight differential pathway enrichment across neutrophil subpopulations.

### Statistical analysis

All statistical parameters, including sample sizes (n), measures of center, dispersion, and precision, are presented in the figures and their corresponding legends. Statistical significance was set at P < 0.05. For comparisons between two groups, two-tailed Student’s t-tests were conducted. For multiple group comparisons, one-way analysis of variance (ANOVA) followed by Tukey’s test was performed. All statistical analyses were carried out using GraphPad Prism software (version 9, GraphPad Software). Each sample was measured once, and all data points were included in the analyses. Statistical tests were conducted with a 95% confidence interval. While normal distribution of data was assumed, formal normality testing was not performed. The experiments were conducted without blinding during data collection and analysis.

## References

1. Mantovani, A., Cassatella, M. A., Costantini, C. & Jaillon, S. Neutrophils in the activation and regulation of innate and adaptive immunity. Nature Reviews Immunology vol. 11 519–531 Preprint at 10.1038/nri3024 (2011).

2. Wculek, S. K. & Malanchi, I. Neutrophils support lung colonization of metastasis-initiating breast cancer cells. Nature 528, 413–417 (2015).

3. McDowell, S. A. C. et al. Neutrophil oxidative stress mediates obesity-associated vascular dysfunction and metastatic transmigration. Nat Cancer 2, 545–562 (2021).

4. Jaillon, S. et al. Neutrophil diversity and plasticity in tumour progression and therapy. Nature Reviews Cancer vol. 20 485–503 Preprint at 10.1038/s41568-020-0281-y (2020).

5. Maas, R. R. et al. The local microenvironment drives activation of neutrophils in human brain tumors. Cell 186, 4546–4566.e27 (2023).

6. Hedrick, C. C. & Malanchi, I. Neutrophils in cancer: heterogeneous and multifaceted. Nature Reviews Immunology vol. 22 173–187 Preprint at 10.1038/s41577-021-00571-6 (2022).

7. Gong, Z., et al. Immunosuppressive reprogramming of neutrophils by lung mesenchymal cells promotes breast cancer metastasis. Sci Immunol 8, (2023).

8. Quail, D. F. et al. Obesity alters the lung myeloid cell landscape to enhance breast cancer metastasis through IL5 and GM-CSF. Nat Cell Biol 19, 974–987 (2017).

9. Palomino-Segura, M., Sicilia, J., Ballesteros, I. & Hidalgo, A. Strategies of neutrophil diversification. Nat Immunol 24, 575–584 (2023).

10. McFarlane, A. J., Fercoq, F., Coffelt, S. B. & Carlin, L. M. Neutrophil dynamics in the tumor microenvironment. Journal of Clinical Investigation vol. 131 Preprint at 10.1172/JCI143759 (2021).

11. Lavillegrand, J. R. et al. Alternating high-fat diet enhances atherosclerosis by neutrophil reprogramming. Nature (2024) doi:10.1038/s41586-024-07693-6.

12. Granot, Z. et al. Tumor entrained neutrophils inhibit seeding in the premetastatic lung. Cancer Cell 20, 300–314 (2011).

13. Wu, Y. et al. Neutrophil profiling illuminates anti-tumor antigen-presenting potency. Cell 187, 1422–1439.e24 (2024).

14. Khan, N. et al. β-Glucan reprograms neutrophils to promote disease tolerance against influenza A virus. Nat Immunol 26, 174–187 (2025).

15. Kalafati, L. et al. Innate Immune Training of Granulopoiesis Promotes Anti-tumor Activity. Cell 183, 771–785.e12 (2020).

16. Ballesteros, I. et al. Co-option of Neutrophil Fates by Tissue Environments. Cell 183, 1282–1297.e18 (2020).

17. Gee, M. H. & Albertine, K. H. NEUTROPHIL-ENDOTHELIAL CELL INTERACTIONS IN THE LUNG. Annu. Rev. Physioi vol. 55 www.annualreviews.org (1993).

18. Hirsch, F. R. et al. Lung cancer: current therapies and new targeted treatments. Lancet 389, 299–311 (2017).

19. Nguyen, D. X., Bos, P. D. & Massagué, J. Metastasis: from dissemination to organ-specific colonization. Nat Rev Cancer 9, 274–284 (2009).

20. Lee, A. H. & Dixit, V. D. Dietary Regulation of Immunity. Immunity vol. 53 510–523 Preprint at 10.1016/j.immuni.2020.08.013 (2020).

21. Mittelman, S. D. The Role of Diet in Cancer Prevention and Chemotherapy Efficacy. Annual Review of Nutrition vol. 40 273–297 Preprint at 10.1146/annurev-nutr-013120-041149 (2020).

22. Radzikowska, U. et al. The influence of dietary fatty acids on immune responses. Nutrients vol. 11 Preprint at 10.3390/nu11122990 (2019).

23. Prendeville, H. & Lynch, L. Diet, lipids, and antitumor immunity. Cellular and Molecular Immunology vol. 19 432–444 Preprint at 10.1038/s41423-021-00781-x (2022).

24. Kanarek, N., Petrova, B. & Sabatini, D. M. Dietary modifications for enhanced cancer therapy. Nature vol. 579 507–517 Preprint at 10.1038/s41586-020-2124-0 (2020).

25. Zhu, H. et al. Ketogenic diet for human diseases: the underlying mechanisms and potential for clinical implementations. Signal Transduction and Targeted Therapy vol. 7 Preprint at 10.1038/s41392-021-00831-w (2022).

26. Steck, S. E. & Murphy, E. A. Dietary patterns and cancer risk. Nature Reviews Cancer vol. 20 125–138 Preprint at 10.1038/s41568-019-0227-4 (2020).

27. Bojková, B., Winklewski, P. J. & Wszedybyl-Winklewska, M. Dietary fat and cancer— which is good, which is bad, and the body of evidence. International Journal of Molecular Sciences vol. 21 1–56 Preprint at 10.3390/ijms21114114 (2020).

28. Christ, A., Lauterbach, M. & Latz, E. Western Diet and the Immune System: An Inflammatory Connection. Immunity 51, 794–811 (2019).

29. Belabed, M. et al. Cholesterol mobilization regulates dendritic cell maturation and the immunogenic response to cancer. Nat Immunol (2025) doi:10.1038/s41590-024-02065-8.

30. Wang, T. et al. Influenza-trained mucosal-resident alveolar macrophages confer long-term antitumor immunity in the lungs. Nat Immunol (2023) doi:10.1038/s41590-023-01428-x.

31. Mizutani, T. et al. Conditional IFNAR1 ablation reveals distinct requirements of Type I IFN signaling for NK cell maturation and tumor surveillance. Oncoimmunology 1, 1027–1037 (2012).

32. Linde, I. L. et al. Neutrophil-activating therapy for the treatment of cancer. Cancer Cell 41, 356–372.e10 (2023).

33. Anderson, K. G. et al. Intravascular staining for discrimination of vascular and tissue leukocytes. Nat Protoc 9, 209–222 (2014).

34. Monsel, A. et al. Analysis of autofluorescence in polymorphonuclear neutrophils: A new tool for early infection diagnosis. PLoS One 9, (2014).

35. Mallick, R. & Duttaroy, A. K. Modulation of endothelium function by fatty acids. Molecular and Cellular Biochemistry vol. 477 15–38 Preprint at 10.1007/s11010-021-04260-9 (2022).

36. Jiang, H. et al. Palmitic acid promotes endothelial progenitor cells apoptosis via p38 and JNK mitogen-activated protein kinase pathways. Atherosclerosis 210, 71–77 (2010).

37. Jiang, H. et al. Mechanisms of Oxidized LDL-Mediated Endothelial Dysfunction and Its Consequences for the Development of Atherosclerosis. Frontiers in Cardiovascular Medicine vol. 9 Preprint at 10.3389/fcvm.2022.925923 (2022).

38. Higashi, Y. Endothelial Function in Dyslipidemia: Roles of LDL-Cholesterol, HDL-Cholesterol and Triglycerides. Cells vol. 12 Preprint at 10.3390/cells12091293 (2023).

39. Bae, J.-H., et al. Postprandial Hypertriglyceridemia Impairs Endothelial Function by Enhanced Oxidant Stress. Atherosclerosis vol. 155 www.elsevier.com (2001).

40. Yunoki, K. et al. Impact of hypertriglyceridemia on endothelial dysfunction during statin ± ezetimibe therapy in patients with coronary heart disease. American Journal of Cardiology 108, 333–339 (2011).

41. Kita, T. et al. Role of oxidized LDL in atherosclerosis. in Annals of the New York Academy of Sciences vol. 947 199–206 (New York Academy of Sciences, 2001).

42. Bui, T. M., Wiesolek, H. L. & Sumagin, R. ICAM-1: A master regulator of cellular responses in inflammation, injury resolution, and tumorigenesis. Journal of Leukocyte Biology vol. 108 787–799 Preprint at 10.1002/JLB.2MR0220-549R (2020).

43. Yang, L. et al. ICAM-1 regulates neutrophil adhesion and transcellular migration of TNF-α-activated vascular endothelium under flow. Blood 106, 584–592 (2005).

44. Quake, S. R. Tabula Sapiens reveals transcription factor expression, senescence effects, and sex-specific features in cell types from 28 human organs and tissues. Preprint at 10.1101/2024.12.03.626516 (2024).

45. Goveia, J. et al. An Integrated Gene Expression Landscape Profiling Approach to Identify Lung Tumor Endothelial Cell Heterogeneity and Angiogenic Candidates. Cancer Cell 37, 21–36.e13 (2020).

46. Salcher, S. et al. High-resolution single-cell atlas reveals diversity and plasticity of tissue-resident neutrophils in non-small cell lung cancer. Cancer Cell 40, 1503–1520.e8 (2022).

47. Pillay, J. et al. Effect of the CXCR4 antagonist plerixafor on endogenous neutrophil dynamics in the bone marrow, lung and spleen. J Leukoc Biol 107, 1175–1185 (2020).

48. Devi, S. et al. Neutrophil mobilization via plerixaformediated CXCR4 inhibition arises from lung demargination and blockade of neutrophil homing to the bone marrow. Journal of Experimental Medicine 210, 2321–2336 (2013).

49. Xie, H. J. et al. Effect of body mass index on survival of patients with stage I non-small cell lung cancer. Chin J Cancer 36, (2017).

50. Shi, D. et al. An isocaloric moderately high-fat diet extends lifespan in male rats and Drosophila. Cell Metab 33, 581–597.e9 (2021).

51. Augustin, H. G. & Koh, G. Y. A systems view of the vascular endothelium in health and disease. Cell vol. 187 4833–4858 Preprint at 10.1016/j.cell.2024.07.012 (2024).

52. Lin, X. et al. Blood lipids profile and lung cancer risk in a meta-analysis of prospective cohort studies. J Clin Lipidol 11, 1073–1081 (2017).

53. Karayama, M. et al. Increased serum cholesterol and long-chain fatty acid levels are associated with the efficacy of nivolumab in patients with non-small cell lung cancer. Cancer Immunology, Immunotherapy 71, 203–217 (2022).

54. Matyash, V., Liebisch, G., Kurzchalia, T. V., Shevchenko, A. & Schwudke, D. Lipid extraction by methyl-terf-butyl ether for high-throughput lipidomics. in Journal of Lipid Research vol. 49 1137–1146 (2008).

