## Supplemental data for "Fat-rich diet reprograms intrapulmonary neutrophils to boost tissue-specific antitumor immunity"

### Supplementary figures and legends

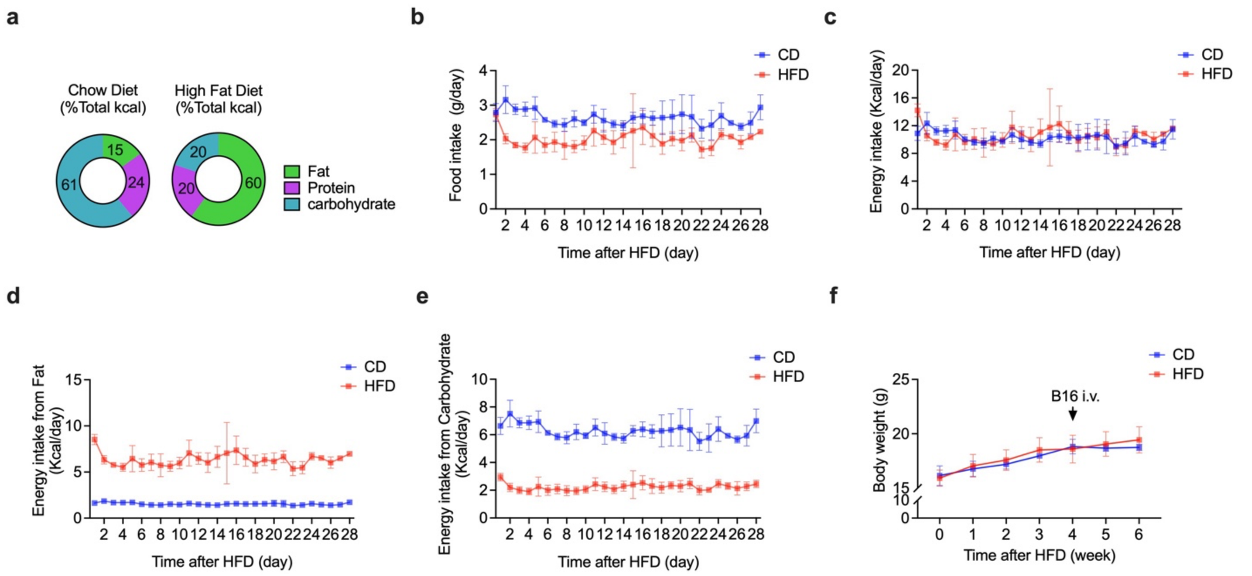


**Extended data figure 1. Short-term HFD is isocaloric**

(**a**) Percentages of calories derived from fat, protein, and carbohydrates in CD and HFD.

(**b**) Daily food intake in mice. In (**b-e**), n = 5 cages of mice on days 1 to 23, n = 4 cages of mice on day 24, and n = 2 cages of mice on days 25 to 28.

(**c**) Calculated daily energy intake in mice.

(**d**) Daily calorie intake from fat.

(**e**) Daily calorie intake from carbohydrate.

(**f**) Kinetics of body weight during the period of HFD versus CD feeding. n = 4 mice in CD group, and n = 5 mice in HFD group.

Data are representative of three independent experiments. Graphs with error bars are presented as mean ± s.d. Two-tailed Student's t-test was used for comparison between two-groups.

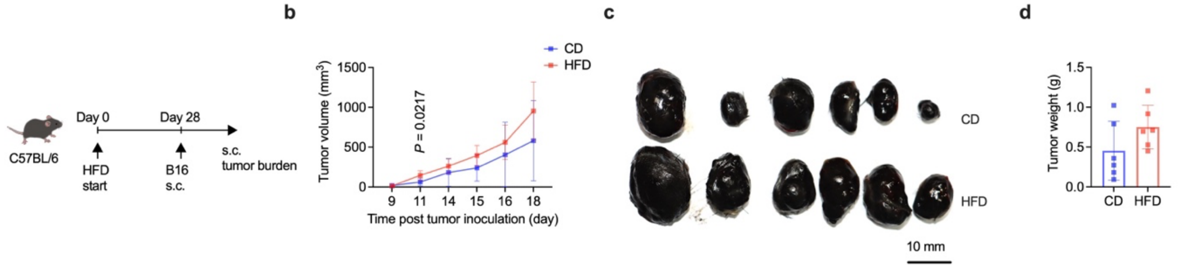


**Extended data figure 2. HFD does not alter systemic antitumor immunity**

(**a**) Schema of subcutaneous (s.c.) B16 melanoma tumor model in CD and HFD mice.

(**b**) Kinetic changes in subcutaneous B16 tumor size at the injection site. In (**b-d**), n = 6 mice per group.

(**c**,**d**) Macroscopic appearance (**c**) and weight (**d**) of subcutaneous B16 tumors at the experimental endpoint.

Data are representative of two independent experiments. Graphs with error bars are presented as mean ± s.d. Two-tailed Student's t-test was used for comparison between two-groups.

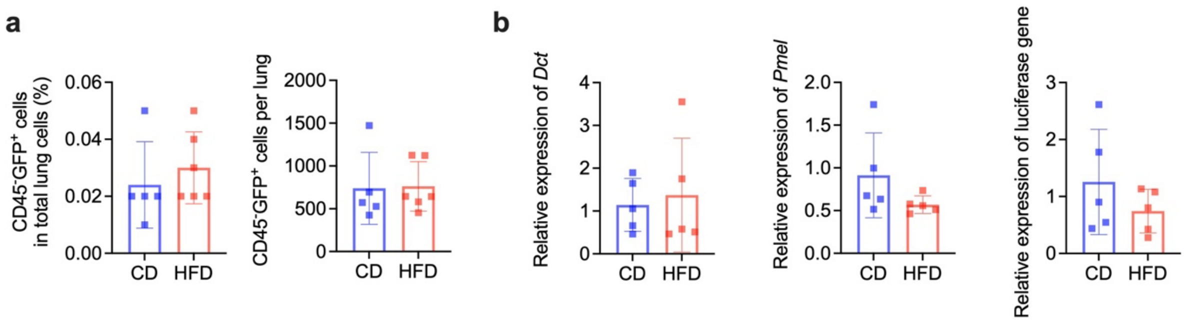


**Extended data figure 3. HFD does not alter initial inoculation of B16 tumor cells in lung tissues**

(**a**) Frequency and absolute number of GFP-expressing B16-luc melanoma cells detected by flow cytometry in the lungs of CD or HFD mice at 24 hours post intravenous tumor inoculation. n = 5 mice in CD group, n = 6 mice in HFD group.

(**b**) RT-PCR-based quantification of B16-luc cells in the lungs of CD or HFD mice at 24 hours post tumor cell inoculation. n = 5 mice per group.

Data are representative of two independent experiments. Bar graphs are presented as mean ± s.d. Two-tailed Student's t-test was used for comparison between two-groups.

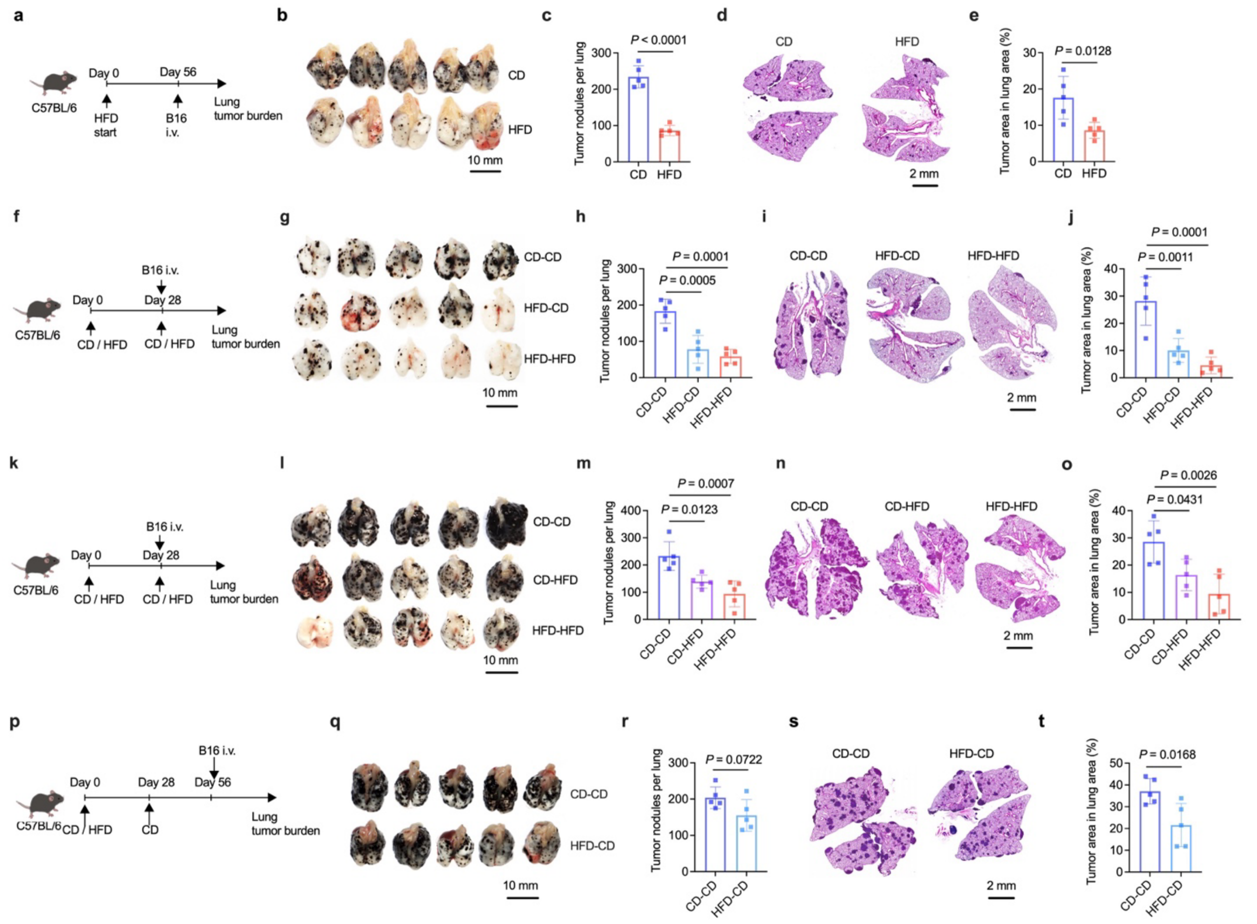


**Extended data figure 4. Pulmonary antitumor immunity is induced by HFD feeding before and after tumor inoculation**

(**a**) Schema of a 56-day HFD feeding schedule in mice followed by i.v. injection of B16 melanoma cells.

(**b**,**c**) Representative macroscopic lung images (**b**) and quantification of visible tumor nodules (**c**) on the surface of lung lobes. n = 5 mice per group.

(**d**,**e**) Representative lung histopathology images (**d**) and percentage of lung area occupied by tumor lesions (**e**) based on lung histopathological analysis in mice shown in (**b**).

(**f**) Schema of B16 melanoma cell i.v. inoculation in mice fed with HFD for an initial 28 days followed by continued HFD feeding after tumor inoculation or switching to CD immediately after tumor inoculation.

(**g**,**h**) Representative macroscopic lung images (**g**) and quantification of visible tumor nodules (**h**) on the surface of lung lobes. n = 5 mice per group.

(**i**,**j**) Representative lung histopathology images (**i**) and percentage of lung area occupied by tumor lesions (**j**) based on lung histopathological analysis in mice shown in (**g**).
(**k**) Schema of B16 melanoma cell i.v. inoculation in mice with HFD started 28 days before or immediately after tumor inoculation.
(**l**,**m**) Representative macroscopic lung images (**l**) and quantification of visible tumor nodules (**m**) on the surface of lung lobes. n = 5 mice per group.
(**n**,**o**) Representative lung histopathology images (**n**) and percentage of lung area occupied by tumor lesions (**o**) based on lung histopathological analysis in mice shown in (**l**).
(**p**) Schema of mice fed with HFD for 28 days and then switched to CD for another 28 days, followed by B16 melanoma cell i.v. inoculation.
(**q**,**r**) Representative macroscopic lung images (**q**) and quantification of visible tumor nodules (**r**) on the surface of lung lobes. n = 5 mice per group.
(**s**,**t**) Representative lung histopathology images (**s**) and percentage of lung area occupied by tumor lesions (**t**) based on lung histopathological analysis in mice shown in (**q**).
Data are representative of two (**g**-**t**) or three (**b**-**e**) independent experiments. Bar graphs are presented as mean ± s.d. Two-tailed Student's t-test was used for comparison between two-groups. One-way ANOVA followed by a Tukey test was performed to compare more than two groups.

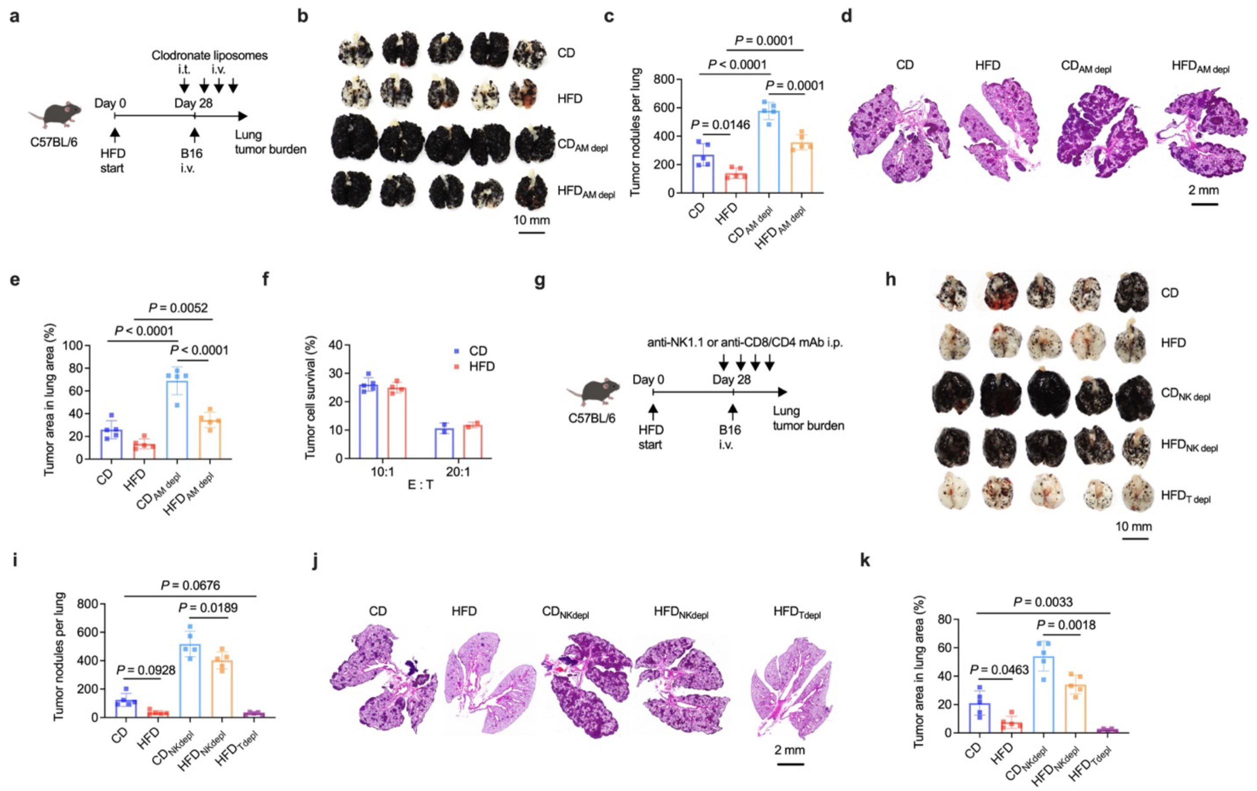


**Extended data figure 5. HFD-induced antitumor immunity is independent of AMs, T cells or NK cells**

(**a**) Schema of AMs depletion in CD and HFD mice followed by i.v. inoculation of B16 melanoma cells.

(**b**,**c**) Representative macroscopic lung images (**b**) and quantification of visible tumor nodules (**c**) on the surface of lung lobes. n = 5 mice per group.

(**d**,**e**) Representative lung histopathology images (**d**) and percentage of lung area occupied by tumor lesions (**e**) based on lung histopathological analysis in mice shown in (**b**).

(**f**) Survival of B16 melanoma cells cocultured with CD or HFD AMs.

(**g**) Schema of continuous depletion of NK or T cells in CD and HFD mice immediately before and after i.v. inoculation of B16 melanoma cells.

(**h**,**i**) Representative macroscopic lung images (**h**) and quantification of visible tumor nodules (**i**) on the surface of lung lobes. n = 5 mice per group.

(**j**,**k**) Representative lung histopathology images (**j**) and percentage of lung area occupied by tumor lesions (**k**) based on lung histopathological analysis in mice shown in (**h**).

Data are representative of two independent experiments. Bar graphs are presented as mean ± s.d. Two-tailed Student's t-test was used for comparison between two-groups. One-way ANOVA followed by a Tukey test was performed to compare more than two groups.

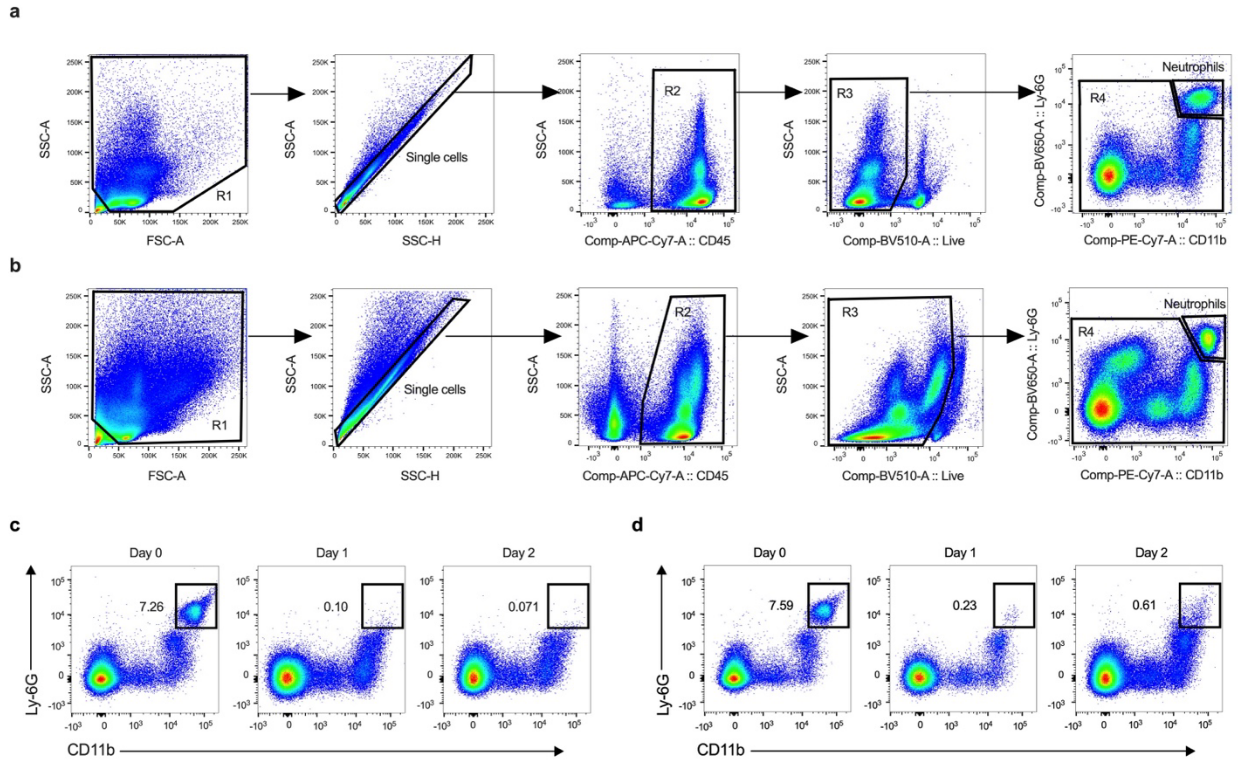


**Extended data figure 6. Flow cytometry gating strategy and depletion of neutrophils *in vivo***

(**a**,**b**) Representative flow cytometry dot plots showing gating strategy for identification of neutrophils in PB (**a**) and lung tissues (**b**).

(**c**,**d**) Representative flow cytometry dot plots showing depletion of neutrophils in PB before (day 0) and after *in vivo* administration of one dose of anti-Ly6G monoclonal antibody in wild type mice (**c**) or diphtheria toxin (DT) in Ly6G-DTR mice (**d**).

Data are representative of two independent experiments. Numbers beside gates represent percentages in parental gates.

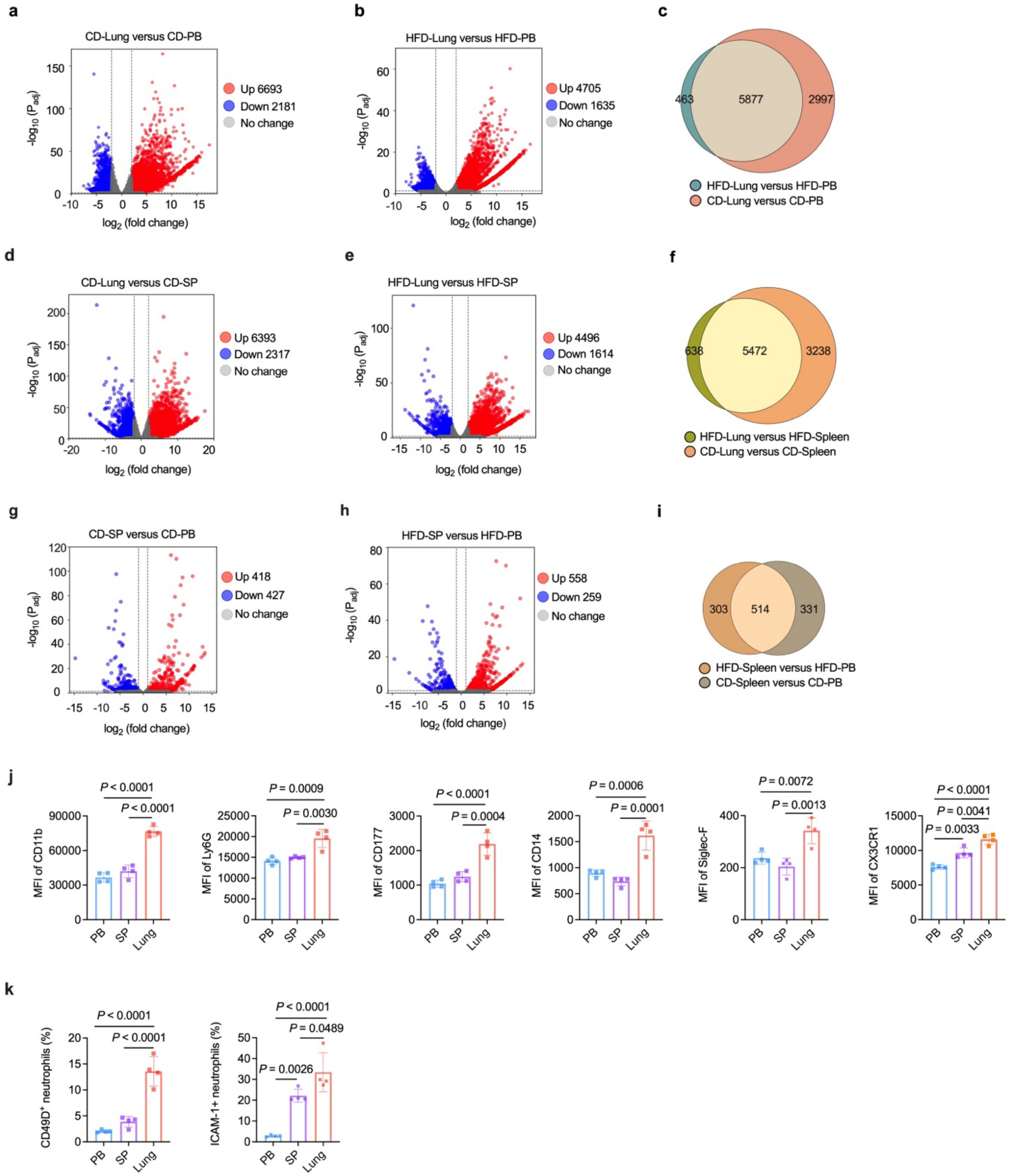


**Extended data figure 7. Lung neutrophils show tissue-specific transcriptional profiles and phenotype**

(**a**,**b**) Volcano plots showing transcriptional differences between lung and peripheral blood (PB) neutrophils in CD (**a**) and HFD (**b**) mice at 28 days after the initiation of HFD. (DESeq2, $P < 0.05$, $|\log2(\text{fold change})| > 2$).

(**c**) Venn diagram showing overlapping of DEGs in a with those b.

(**d**,**e**) Volcano plots showing transcriptional differences between lung and splenic (SP) neutrophils in CD (**d**) and HFD (**e**) mice (DESeq2, $P < 0.05$, $|\log2(\text{fold change})| > 2$).

(**f**) Venn diagram showing overlapping of DEGs in d with those e.

(**g**,**h**) Volcano plots showing transcriptional differences between SP and PB neutrophils in CD (**g**) and HFD (**h**) mice (DESeq2, $P < 0.05$, $|\log2(\text{fold change})| > 1$).

(**i**) Venn diagram showing overlapping of DEGs in (**g**) with those (**h**).

(**j**) Median fluorescence intensity (MFI) of surface markers on PB, SP and lung neutrophils in CD mice.

(**k**) Frequencies of CD49D$^+$ neutrophils and ICAM-1$^+$ neutrophils in total neutrophils from PB, SP or lung of CD mice.

Data in (**j**) and (**k**) are representative of two independent experiments, and data in (**a**-**i**) are from one experiment. Bar graphs are presented as mean ± s.d. In (**a**-**i**), n = 4 in CD-PB and CD lung groups, n = 3 in CD-SP and HFD-lung groups, and n = 2 in HFD-PB and HFD-SP groups. One-way ANOVA followed by a Tukey test was performed to compare more than two groups.

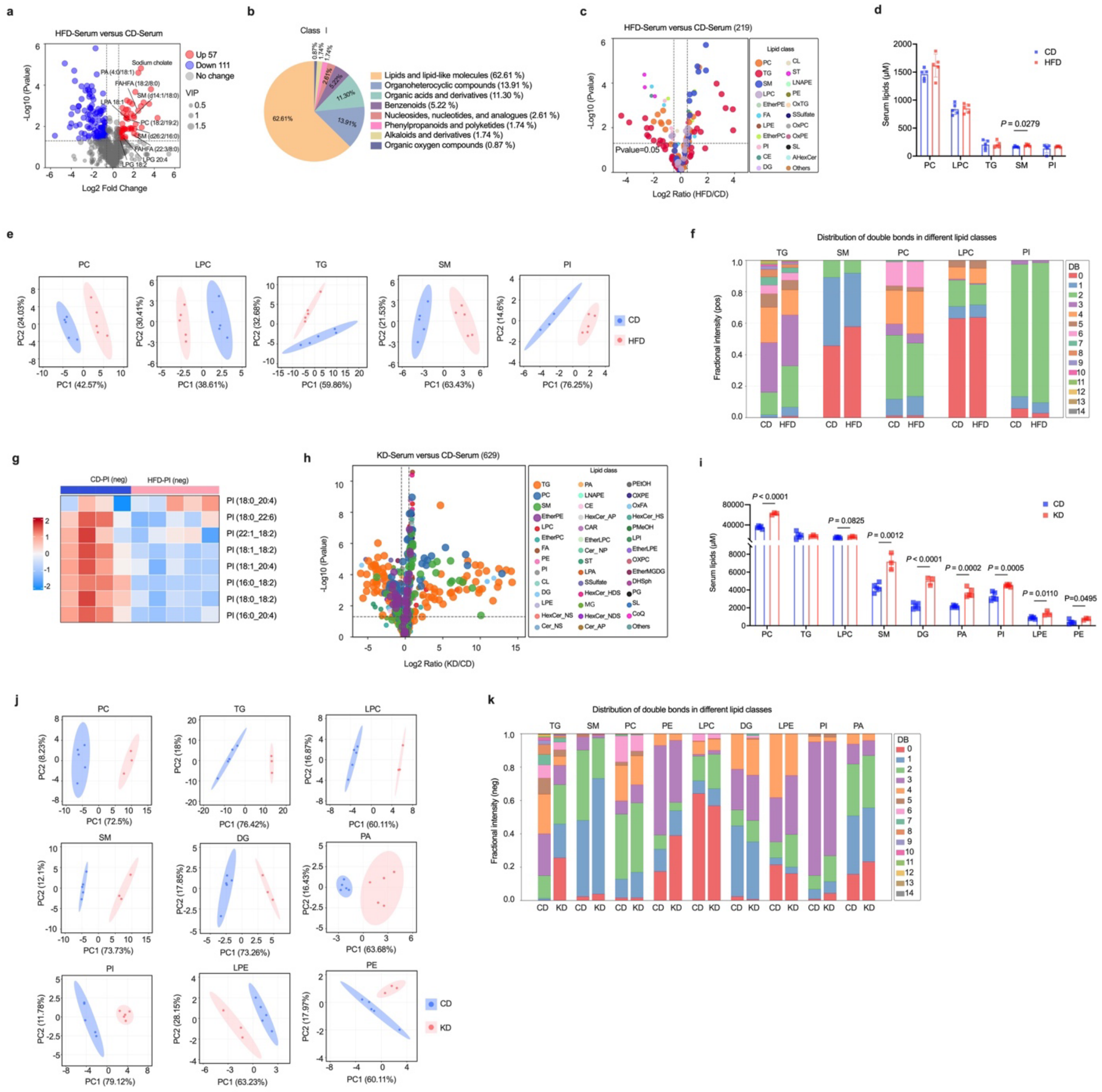


**Extended data figure 8. FRD alters serum lipid contents**

(**a**) Volcano plot showing differential metabolite abundances in serum between HFD and CD mice, analyzed by positive and negative ion mode mass spectrometry. Red and blue dots indicate significantly upregulated and downregulated metabolites, respectively ($P < 0.05$, $|\log2(\text{fold change})| > 0.5$). Dot size corresponds to variable importance in projection (VIP) scores.

(**b**) Pie chart showing the distribution of differential serum metabolites across Class 1 metabolite categories in HFD compared to CD mice.

(**c**) Volcano plot of differentially abundant lipid species in serum of HFD versus CD mice, determined by LC-MS/MS in positive and negative ion modes.

(**d**) Quantification of major lipid classes in serum of CD and HFD mice by LC-MS analysis.

(**e**) PCA plot showing distinct clustering patterns of serum lipidome profiles between CD and HFD mice.

(**f**) Stacked bar plots showing the fractional distribution of serum lipids across lipid classes in HFD versus CD mice, stratified by total fatty acyl-side chain unsaturation. Triglycerides (TG), sphingomyelin (SM), phosphatidylcholines (PC), and lyso-phosphatidylcholine (LPC) were detected by positive-mode LC-MS/MS. Phosphatidylinositol (PI) were detected by negative-mode analysis. DB (0-14) indicates the total number of double bonds in fatty acyl-side chains.

(**g**) Heatmap of significantly altered phosphatidylinositols (PIs) in serum between CD and HFD mice, identified by negative-mode LC-MS analysis ($P < 0.05$).

(**h**) Volcano plot of differentially abundant lipid species in serum of KD versus CD mice, determined by LC-MS/MS in positive and negative ion modes.

(**i**) Quantification of major lipid classes in serum of CD and KD mice by LC-MS analysis.

(**j**) PCA plot showing distinct clustering patterns of serum lipidome profiles between CD and KD mice.

(**k**) Stacked bar plots showing the fractional distribution of serum lipids across lipid classes in KD versus CD mice, stratified by total fatty acyl-side chain unsaturation. Triglycerides (TG), sphingomyelin (SM), phosphatidylcholines (PC), phosphatidylethanolamine (PE), lyso-phosphatidylcholine (LPC), diglyceride (DG), and lyso-phosphatidylethanolamine (LPE) were detected by positive-mode LC-MS/MS. Phosphatidic acid (PA) and phosphatidylinositol (PI) were detected by negative-mode analysis. DB represents the total number of double bonds in fatty acyl-side chains.

Data in are from one experiment. In (**a**,**b**), n = 3 mice per group. In (**c**-**g**), n = 5 mice per group for positive ion mode analysis, n = 4 mice in CD group and n = 5 in HFD group for negative ion mode analysis. In (**h-k**), n = 5 mice in CD group and n = 3 in KD group for positive ion mode analysis, n = 5 mice per group for negative ion mode analysis. Bar graphs in (**d**,**i**) are presented as mean ± s.d.

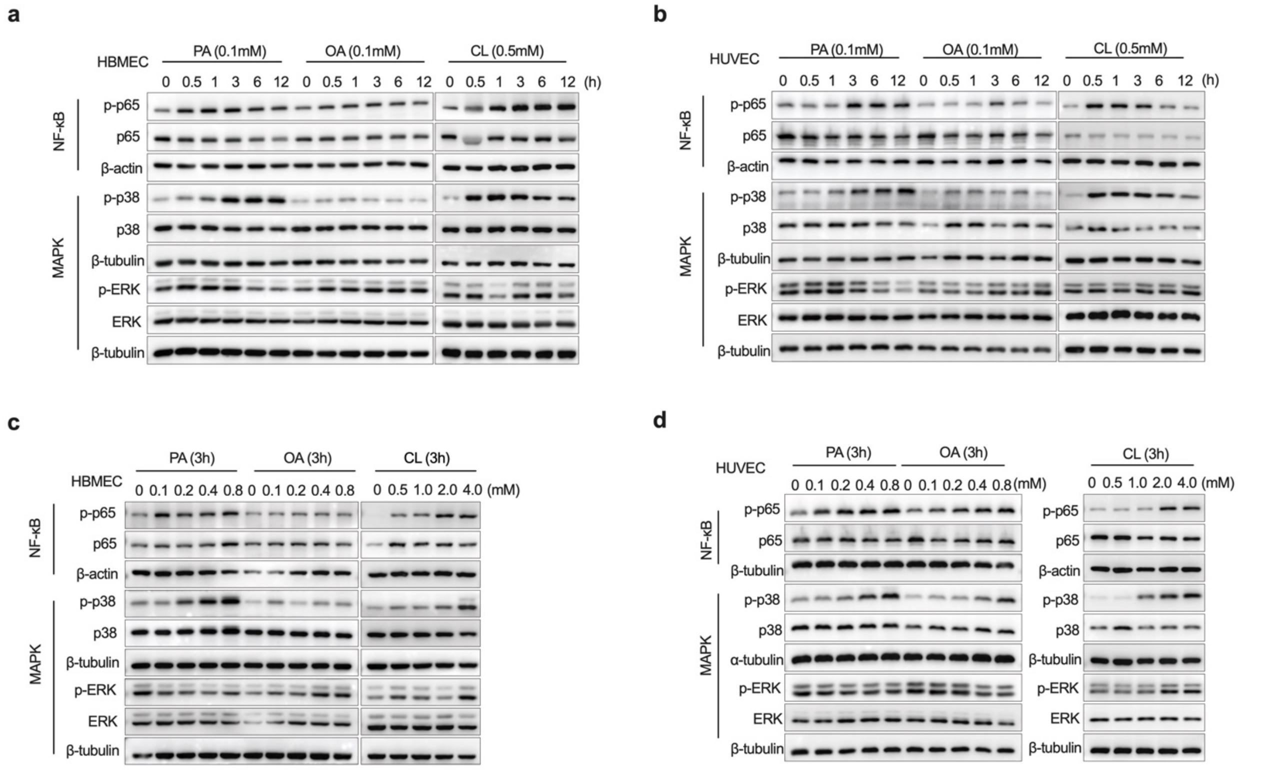


**Extended data figure 9. Saturated and mono-unsaturated fatty acids and cholesterol activate LVECs *in vitro***

(**a**,**b**) Representative immunoblots showing time-dependent activation of signaling pathways in HBMEC (**a**) and HUVEC (**b**) treated with palmitic acid (PA, 0.1 mM), oleic acid (OA, 0.1 mM), or cholesterol (CL, 0.5 mM) for indicated time points (0-12 h). β-actin and tubulin serve as loading controls.

(**c**,**d**) Representative immunoblots showing dose-dependent activation of signaling pathways in HBMEC (**c**) and HUVEC (**d**) exposed to increasing concentrations of PA (0-0.8 mM), OA (0-0.8 mM), or CL (0-4.0 mM) for 3 h. β-actin and tubulin serve as loading controls.

Data are representatives of two independent experiments.
